# UnMICST: Deep learning with real augmentation for robust segmentation of highly multiplexed images of human tissues

**DOI:** 10.1101/2021.04.02.438285

**Authors:** Clarence Yapp, Edward Novikov, Won-Dong Jang, Tuulia Vallius, Yu-An Chen, Marcelo Cicconet, Zoltan Maliga, Connor A. Jacobson, Donglai Wei, Sandro Santagata, Hanspeter Pfister, Peter K. Sorger

**Affiliations:** Laboratory of Systems Pharmacology, Harvard Medical School, Boston, MA 02115; Image and Data Analysis Core, Harvard Medical School, Boston, MA 02115 Human Tumor Atlas Network; School of Engineering and Applied Sciences, Harvard University, Cambridge, MA, 02138; Ludwig Center for Cancer Research at Harvard, Harvard Medical School, Boston, MA, 02115; Department of Pathology, Brigham and Women’s Hospital, Harvard Medical School, Boston, MA, 02115; Department of Systems Biology, Harvard Medical School, Boston, MA, 02115

## Abstract

Newly developed technologies have made it feasible to routinely collect highly multiplexed (20-60 channel) images at subcellular resolution from human tissues for research and diagnostic purposes. Extracting single cell data from such images requires efficient and accurate image segmentation, a challenging problem that has recently benefited from the use of deep learning. In this paper, we demonstrate two approaches to improving tissue segmentation that are applicable to multiple deep learning frameworks. The first uses “real augmentations” that comprise defocused and saturated image data collected on the same instruments as the actual data; using real augmentation improves model accuracy to a significantly greater degree than computational augmentation (Gaussian blurring). The second involves imaging the nuclear envelope to better identify nuclear outlines. The two approaches cumulatively and substantially improve segmentation on a wide range of tissue types and provide a set of improved segmentation models. We speculate that the use of real augmentations may have applications in image processing outside of microscopy.

## INTRODUCTION

Optical microscopy is a method of long-standing importance in biology that is changing rapidly due to the introduction of new methods for highly-multiplexed imaging of normal and diseased tissues (**Supplementary Table 1**)^1–5^. High-plex imaging (sometimes called spatial proteomics) yields subcellular resolution data on the abundance of 20-60 antigens, which is sufficient to identify cell types, measure cell states (quiescent, proliferating, dying, etc.) and interrogate cell signaling pathways. High plex imaging also reveals the morphologies and positions of acellular structures essential for tissue integrity in a preserved 3D environment. The resulting data are substantially more challenging to analyze computationally than images of cultured cells, the primary emphasis of biology-focused machine vision systems to date, but analysis of tissues and similarly complex specimens is benefiting from recent advances in machine learning (deep learning). The different cell types, basement membranes, and connective structures that organize tissues and tumors are present on length scales ranging from subcellular organelles to whole organs (<0.1 to >10^4^ μm). Microscopy using Hematoxylin and Eosin (H&E) complemented by immunohistochemistry^6^ has long played a primary role in the study of tissue architecture^7, 8^ and clinical histopathology remains the primary means by which diseases such as cancer are staged and managed clinically^9^. However, classical histology provides insufficient molecular information to precisely identify cell subtypes, study mechanisms of development, and characterize disease genes. This has motivated development of highly multiplexed (20-60 plex) imaging methods and spatial transcriptomics. These methods differ in resolution, field of view, and multiplicity (plex), but all generate 2D images of tissue sections; in current practice these are usually 5 – 10 μm thick. Developing effective image processing algorithms for the resulting image data is now the primary challenge for molecular profiling of tissues in both research and clinical settings.

**Table 1:** Antibodies used for immunofluorescence staining

| Target | Fluorochrome | Species | Clone | Vendor | Cat. No. | RRID |
| --- | --- | --- | --- | --- | --- | --- |
| DNA | Hoechst 33342 | NA | NA | CST | 4082 | AB_10626776 |
| Lamin B2 | Alexafluor 647 | Rabbit | EPR9701(B) | Abcam | ab200427 | AB_2889288 |
| NUP98 | Alexafluor 647 | Rabbit | C39A3 | CST | 13393 | AB_2728831 |
| Lamin B1 | Alexafluor 488 | Rabbit | EPR8985(B) | Abcam | ab194106 | AB_2728786 |
| Lamin A/C | Alexafluor 488 | Mouse | 4C11 | CST | 8617S | AB_10997529 |
| Lamin B receptor | Alexafluor 488 | Rabbit | E398L | Abcam | ab201532 | AB_2889290 |

When multiplexed images are segmented and quantified, the resulting single cell data are a natural complement to single cell RNA Sequencing (scRNASeq) data, which have had a dramatic impact on our understanding of normal and diseased cells and tissues^10, 11^. Unlike dissociative RNASeq, however, multiplex tissue imaging preserves morphology and spatial information. Single cell analysis of imaging data requires segmentation, a computer vision technique that assigns class labels to an image in a pixel-wise manner to subdivide it; this is followed by quantification of the levels of protein markers on a per-cell or per-organelle basis. Marker quantification typically involves integrating fluorescent signal intensities across a segmentation mask or a shape (usually an annulus) that outlines or is centered in the mask^12^. Extensive work has gone into the development of methods for segmenting metazoan cells grown in culture, but segmentation of tissue images is a more difficult challenge due to cell crowding and the diverse morphologies of different cell types. Recently, segmentation routines that use machine learning have become standard, paralleling the widespread use of convolutional neural networks (CNNs) in image recognition, object detection, and synthetic image generation^13^. Architectures such as ResNet, VGG16, and more recently, UNet and Mask R-CNN^14, 15^ have gained widespread acceptance for their ability to learn millions of parameters and generalize across datasets, as evidenced by excellent performance in a wide range of segmentation competitions, as well as in hackathon challenges^16^ using publicly available image datasets^17, 18^. However, to our knowledge, the relative strengths and weaknesses of different CNN segmentation frameworks has not yet been evaluated directly on tissue images.

In both cultured cells and tissues, localizing nuclei is an optimal starting point for segmenting cells since most cell types have one nucleus (cells undergoing mitosis, muscle and liver cells and osteoclasts are important exceptions), and nuclear stains with high signal-to-background ratios are widely available. The nucleus is generally quite large (5-10 μm) relative to the resolution of wide-field fluorescence microscopes (~0.5 μm for a 0.9 numerical aperture – NA – objective lens), making it easy to detect at multiple magnifications. Nuclei are also often found at the approximate center of a cell. There are possible advantages to using additional markers during image acquisition; for example Schüffler et al.^19^ used multiplexed IMC data and watershed methods for multi-channel segmentation. Methods based on random forests such as Ilastik and Weka^20, 21^ also exploit multiple channels for class-wise pixel classification via an ensemble of decision trees to assign pixel-wise class probabilities in an image. However, random forest models have significantly less capacity for learning than CNNs, which is a substantial disadvantage. The possibility of using CNNs with multi-channel data to enhance nuclei segmentation has not been widely explored.

The accuracy of segmentation algorithms is crucially dependent on the quality of the original images. In practice, many images of human and murine tissues have focus artefacts (blur) and images of some cells are saturated (with intensities above the linear range of the camera). This is particularly true of whole-slide imaging in which up to 1,000 sequentially acquired image tiles are used to create mosaic images of specimens as large as several square centimeters. Whole slide imaging is a diagnostic necessity^22^ and essential to achieve sufficient power for rigorous spatial analysis ^23^. However, many recent papers addressing the segmentation of tissue images restrict their analysis to the clearest in-focus fields. This is logical because, in the setting of supervised learning, it is easier to obtain training data and establish a ground-truth when images are clear and inter-observer agreement is high. In practice however, all microscopy images of tissue specimens have issues with focus: the depth of field of objective lenses capable of high resolution imaging (high NA lenses) is typically less than the thickness of the specimen so that objects above and below the plane of optimal focus are blurred. Images of human biopsy specimens are particularly subject to blur and saturation artefacts because the tissue sections are not always uniformly co-planar with the cover slip. Since most research on human tissues is incidental to diagnosis or treatment, it is rarely possible to reject problematic specimens outright. Moreover, reimaging of previously analyzed tissue sections is rarely possible due to tissue disintegration. Thus, image segmentation with real-world data must compensate for common image aberrations.

The most common way to expand training data to account for image artefacts is via augmentation^24^ which involves pre-processing images via random rotation, shearing, flipping, etc. This is designed to prevent algorithms from learning irrelevant aspects of an image, such as orientation. To date, focus artifacts have been tackled using computed Gaussian blur to augment training data^25–27^. However, Gaussian blur is only an approximation of the blurring inherent to any optical imaging system having limited bandpass (that is – any real microscope) plus the effects of refractive index mismatches and light scattering.

In this paper, we investigate ways to maximize the accuracy of image segmentation in multiplexed tissue images containing common imaging artefacts. We generate a set of training and test data with ground-truth annotations via human curation of multiple normal tissues and tumors, and use these data to score segmentation accuracy achieved on three deep learning networks, each of which was independently trained and evaluated: UNet, Mask R-CNN, and Pyramid Scene Parsing Network (PSPNet). The resulting models comprise a family of *Universal Models for Identifying Cells and Segmenting Tissue* (UnMICST) in which each model is based on the same training data but a different class of ML network. Based on our analysis we identify two ways to improve segmentation accuracy for all three networks. The first involves adding images of nuclear envelope staining (NES) to images of nuclear chromatin acquired using DNA-intercalating dyes. The second involves adding “real augmentations”, defined here as intentionally defocused and over-saturated images (collected from the same specimens), to the training data to make models more robust to the types of artefacts encountered in real tissue images. We find that augmentation with real data significantly outperforms conventional Gaussian blur augmentation, offering a statistically significant improvement in model robustness. Across a range of tissue types, improvements from adding NES data and real augmentations are cumulative.

## RESULTS

### Data sets and ground truth annotation of nuclear boundaries

One challenge in supervised machine learning on tissue images is a lack of sufficient freely-available data with ground truth labelling. Experience with natural scene images^14^ has shown that the acquisition of labels can be time consuming and rate limiting^28^. It is also well established that cells in different types of tissue have nuclear morphologies that vary substantially from the spherical and ellipsoidal shape observed in cultured cells^31^. Nuclear pleomorphism (variation in nuclear size and shape) is even used in histopathology to grade cancers^32^. To account for variation in nuclear morphology we generated training, validation, and test datasets from seven different tissue and tumor types (lung adenocarcinoma, non-neoplastic small intestine, normal prostate, colon adenocarcinoma, glioblastoma, non-neoplastic ovary, and tonsil) found in 12 cores from EMIT (Exemplar Microscopy Images of Tissue^33^, RRID: SCR_021052), a tissue microarray assembled from clinical discards. The tissues had cells with nuclear morphologies ranging from mixtures of cells that were large vs. small, round cells vs. narrow, and densely and irregularly packed vs. organized in clusters. A total of ~10,400 nuclei were labelled by a human expert for nuclear contours, centers, and background. In addition, two human experts labelled a second dataset from a whole-slide image of human melanoma^34^ to establish the level of inter-observer agreement and to provide a test data set that was disjoint from the training data.

### Evaluating the performance of ML segmentation algorithms and models

We implemented and then evaluated two semantic and one instance segmentation algorithms that are based on deep learning/CNNs (UNet, PSPNet, and Mask R-CNN, respectively). Semantic segmentation is a coarse-grained ML approach that assigns objects to distinct trained classes, while instance segmentation is fine grained and identifies individual instances of objects. We trained each of these models (UnMICST-U, UnMICST-P, and UnMICST-M, respectively) on manually curated and labelled data from seven distinct tissue types. The models were not combined, but were tested independently in an attempt to determine which network exhibited the best performance.

A variety of metrics have been proposed for evaluating pixel-level and instance-level segmentation. An example of a pixel-level metric is the sweeping intersection over union (IoU) threshold described by Caicedo et al.^16^ and implemented in the widely used COCO dataset, which is based on images of cell lines^30^. The IoU (the Jaccard Index) is calculated by measuring the overlap between the ground truth annotation and the prediction via a ratio of the intersection to the union of pixels in two masks. The greater the IoU, the higher the accuracy, with an ideal value of 1 (although this is very rarely achieved). The (IoU) threshold is evaluated over a range of values from the least stringent, 0.55, to most stringent, 0.8^16^. Unlike a standard pixel accuracy metric (the fraction of pixels in an image that were correctly classified), IoU is not sensitive to class-imbalance. IoU is a particularly relevant measure of segmentation performance for analysis of high-plex images. When masks are used to quantify marker intensities in other channels, we are concerned not only with whether a nucleus is present or not at a particular location but whether the masks are the correct size and shape.

Examples of instance-level metrics are true positives (TP) and true negatives (TN), which classify predicted objects based on whether they overlap by 50% or greater, otherwise they are deemed as false positives (FP) and false negatives (FN). The frequencies of these four states are used to calculate the F1-score and average precision (AP). The F1-score is the weighted average of precision (true positives normalized to predictions) and recall (true positives normalized to ground truth), and AP considers the number of true positives, total number of ground truth and predictions.

A common approach judging the accuracy expected for a supervised learning method is to use multiple human experts to label the same set of data and determine the level of inter-observer agreement (of course, it may ultimately be possible to exceed this level of human performance ^24, 29^). We assessed inter-observer agreement using both the F1-score and sweeping IoU scores with data from whole-side images of human melanoma^34^. For a set of ~4,900 independently annotated nuclear boundaries, two experienced microscopists achieved a mean F1-score of 0.78 (**Supplementary Figure 1**) and an IoU of 60% at a threshold of 0.6. In the discussion we compare these data to values obtained in other recently published papers and address the discrepancy in F1-scores and IoU values.

### Real augmentations increase model robustness to focus artefacts

To study the impact of real and computed augmentations on the performance of segmentation methods, we trained models with different sets of data involving both real and computed augmentations and then tested the data on images that were acquired in focus, out of focus or blurred using a Gaussian kernel. Where dataset sizes were unbalanced, we supplemented such instances with rotation augmentations. We assessed segmentation accuracy quantitatively based on IoU and qualitatively by visual inspection of predicted masks overlaid on image data. Real augmentation involved adding additional empirical, rather than computed, training data having the types of imperfections most commonly encountered in tissue. This was accomplished by positioning the focal plane 3 μm above and below the specimen, resulting in de-focused images. A second set of images was collected at long exposure times, thereby saturating 70-80% of pixels. Because blurred and saturated images were collected sequentially without changing stage positions, it was possible to use the same set of ground truth annotations. For computed augmentations, we convolved a Gaussian kernel with the in-focus images using a range of standard deviations chosen to cover a broad spectrum of experimental cases (**Figure 1a**). In both scenarios, the resulting models were evaluated on a test set prepared in the same way as the training set.

**Figure 1:**
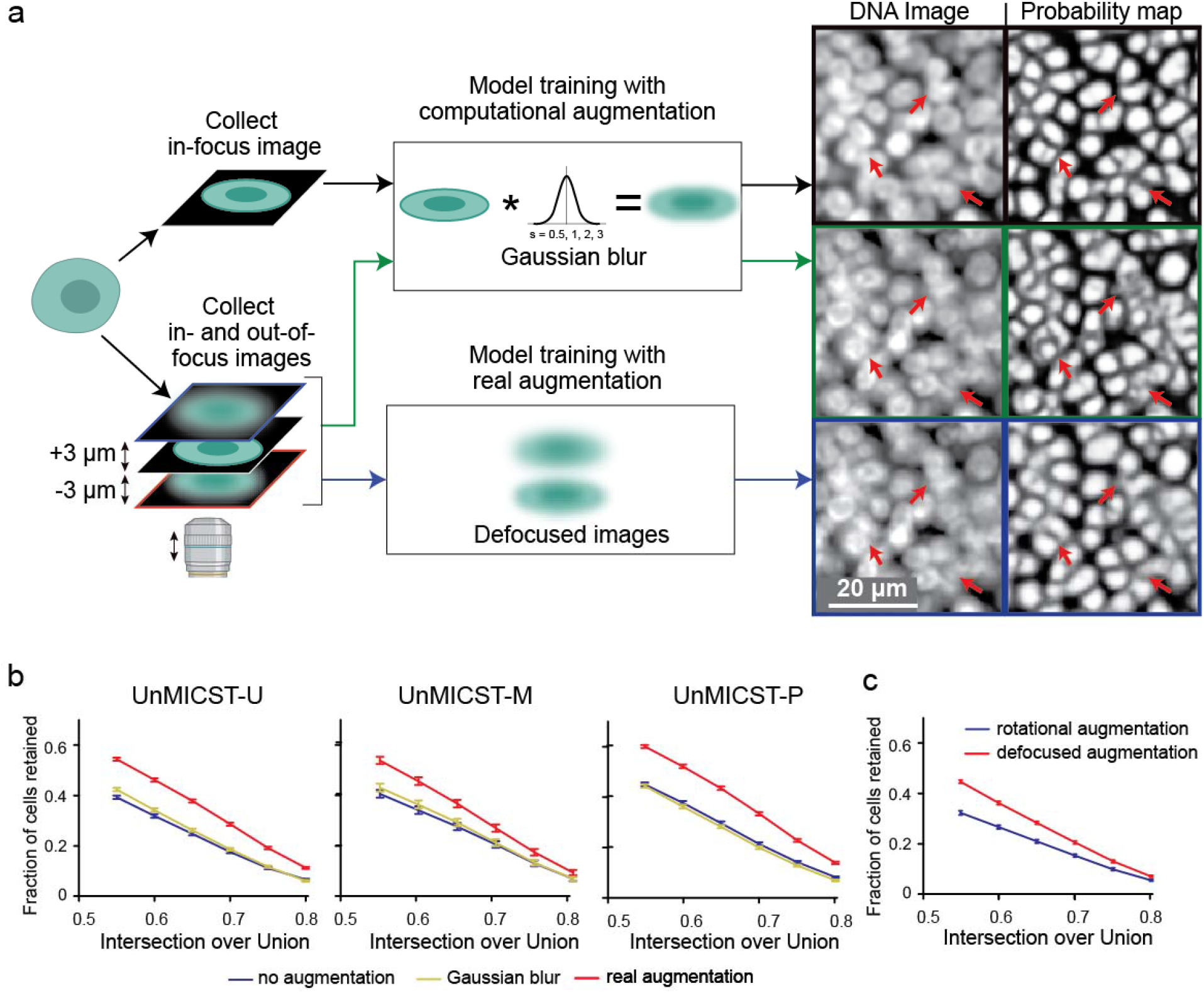
Comparing the use of real augmentations (defocused and overexposed images) and Gaussian blur. **a**) Schematic diagram showing the approach comparing test images on models trained with Gaussian-blurred or defocused image data. Higher contrast probability maps signify more confidence – areas of interest are highlighted with red arrows. Corresponding probability maps indicate a model trained with defocused images performs better on defocused test images than a Gaussian-blurred model. Scale bar denotes 20 micrometers. **b**) Plots show that incorporating real augmentations (red curve) into the training set is statistically significantly superior to training sets with Gaussian blur (yellow curve) and without real augmentations (blue curve) for UnMICST-U, UnMICST-M, and UnMICST-P. Simulating defocused images with Gaussian blur is only marginally better than not augmenting the training data at all. **c**) Comparing UnMICST-U model accuracy when the training dataset size was held constant by replacing defocused augmentations (red curve) with 90 and 180 degree rotations (blue curve).

In an initial set of studies, we found that models created using training data augmented with Gaussian blur performed well on Gaussian blurred test data. However, when evaluated against test data involving defocused and saturated images, we found that Gaussian blur augmentation improved accuracy only slightly relative to baseline models lacking augmentations (**Figure 1b**). In contrast, the use of training data supplemented with real augmentations increased the fraction of cells retained at an IoU threshold of 0.6 by 40-60%. Statistically significant improvement was observed up to an IoU cutoff of 0.8 with all three learning frameworks (UnMICST-U, UnMICST-M, and UnMICST-P models). To perform a balanced comparison, we created two sets of training data having equal numbers of images. The first set contained the original data plus computed 90- and 180-degree rotations, and the second set contained original data plus defocused data collected from above and below the specimen. Again, we found that models trained with real augmentations substantially outperformed rotationally augmented models when tested on defocused test data (**Figure. 1c**). Thus, training any of the three different deep learning architectures with real augmentation generated models that outperformed models with computed augmentation using test data that contained commonly encountered artefacts.

### Addition of NES improves segmentation accuracy

When we stained our TMA panel (the Exemplar Microscopy Images of Tissues and Tumors (EMIT) TMA) we found that antibodies against lamin A and C (**Figure 2a)** (which are different splice forms of *LMNA* gene) stained approximately only half as many nuclei as antibodies against lamin B1 (**Figure 2b**) or lamin B2 (**Figure 2c**) (products of the *LMNB1* and *LMNB2* genes). Staining for the lamin B receptor (**Figure 2e**) exhibited poor image contrast. A pan-tissue survey showed that a mixture of antibodies for nucleoporin NUP98 and lamin B2 conjugated to the same fluorophore (Alexafluor-647) generated nuclear envelope staining (NES) for nearly all nuclei across multiple tissues (**Figure 2f–h**). We judged this to be the optimal antibody cocktail. However, only some cell types, epithelia in colorectal adenocarcinoma for example, exhibited the ring-like structure that is characteristic of nuclear lamina in cultured epithelial cells. The nuclear envelope in immune and other cells has folds and invaginations^35^ and in our data, NES staining could be irregular and diffuse, further emphasizing the difficulty of finding a broadly useful NES stain in tissue.

**Figure 2:**
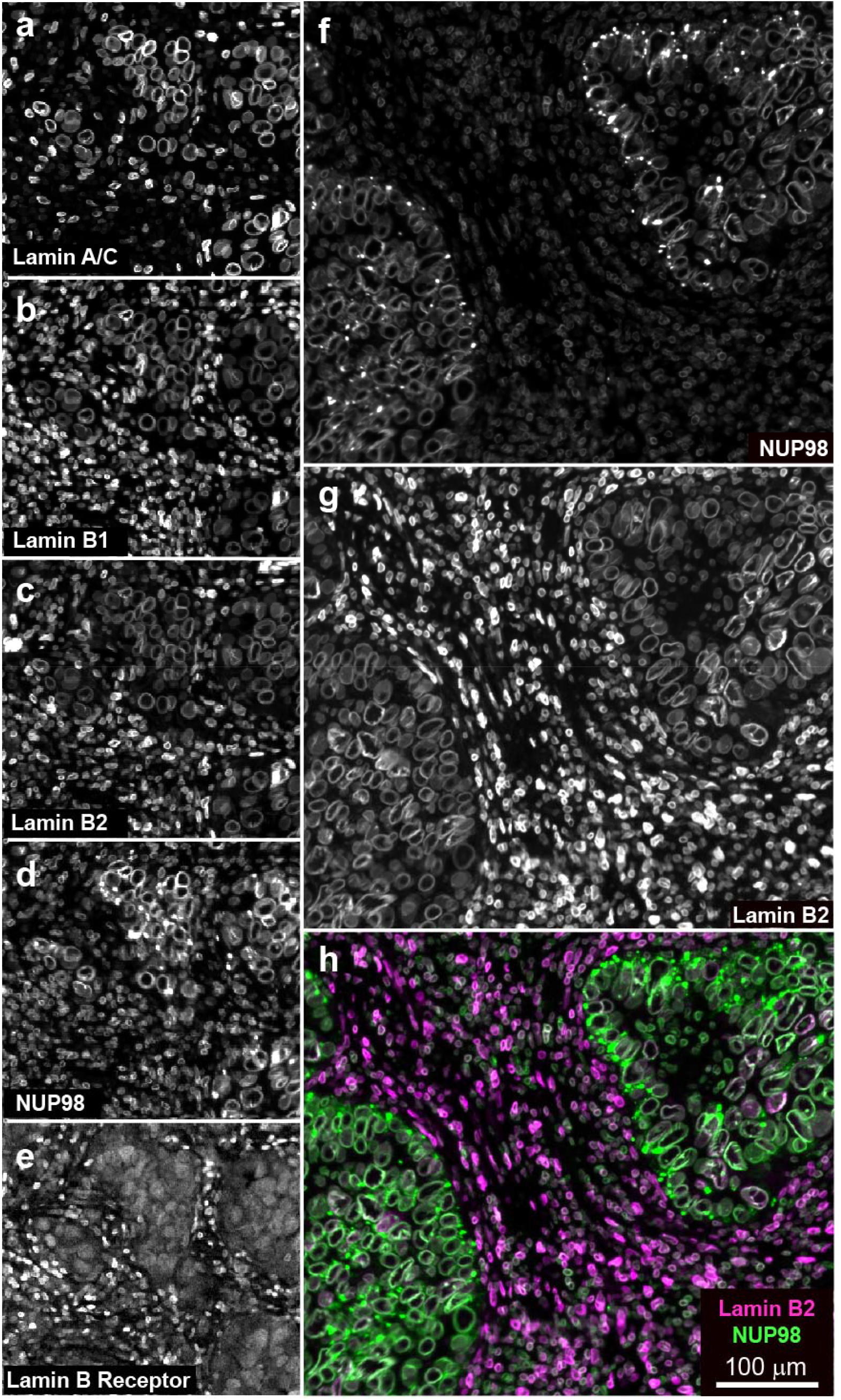
Comparing different nuclear envelope stains in colon adenocarcinoma. **a-e**) Showcasing lamin A/C, lamin B1, lamin B2, NUP98, and the lamin B receptor in the same field of view. Lamin B1 and B2 appear to stain similar proportions of nuclei while lamin A/C stains fewer nuclei. The stain against the lamin B receptor was comparatively weaker. Lamin B2 (**f**) and NUP98 (**g**) are complementary and, when used in combination, maximize the number of cells stained. **h**) Composite of lamin B2 (purple) and NUP98 (green). Scale bar denotes 100 micrometers.

The value of NES images for model performance was assessed quantitatively and qualitatively. In images of colon adenocarcinoma, non-neoplastic small intestine, and tonsil tissue, we found that the addition of NES images resulted in significant improvements in segmentation accuracy based on IoU with all three learning frameworks; improvements in other tissues, such as lung adenocarcinoma, were more modest and sporadic (**Figure 3a**, Lung). For nuclear segmentation of fibroblasts in prostate cancer tissue, UnMICST-U and UnMICST-M models with NES data were no better than models trained on DNA staining alone. Most striking were cases in which NES data slightly decreased performance (UnMICST-P segmentation on prostrate fibroblasts and UnMICST-U segmentation of glioblastoma). Inspection of the UnMICST-P masks suggested that the segmentation of well-separated fibroblast nuclei was already optimal with DNA images alone (~60% of nuclei retained at IoU of 0.6), implying that the addition of NES images afforded little improvement. With UnMICST-U masks in glioblastoma, the problem appeared to involve atypical NES morphology, which is consistent with a high level of nuclear pleomorphism and the presence of “giant cells,” both of which are well-established features of high-grade glioblastoma^36, 37^. We also note that NES data alone was inferior to DNA staining as a sole source of training data and should therefore be used in combination with images of DNA (**Supplementary Figure 2**). Thus, adding NES to training data broadly but not universally improves segmentation accuracy.

**Figure 3:**
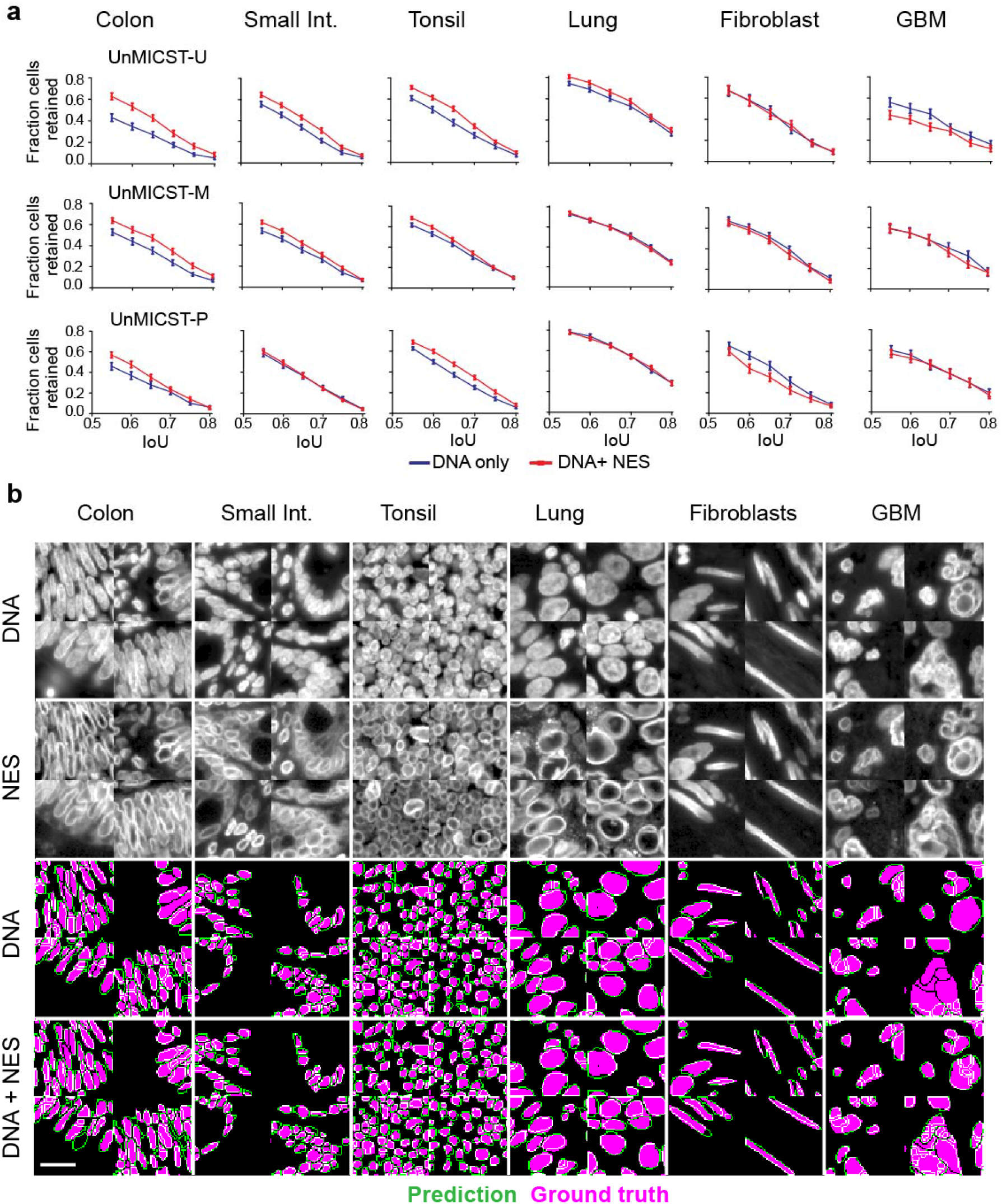
NES with DNA improves nuclear segmentation. NES – nuclear envelope staining. Assessing the addition of NES as a 2^nd^ marker to DNA on segmentation accuracy on a per tissue and per model basis. **a**) Variable IoU plots comparing the DNA-only model (blue curve) and the DNA + NES model (red curve) across frameworks. Adding NES increased accuracy for densely packed nuclei such as colon, small intestine, tonsil, and to some extent, lung tissue. Error bars are standard errors of mean. **b**) Representative grayscale images of tissues stained with DNA and NES comparing their variable morphologies, followed by UnMICST-U mask predictions (green) overlaid onto ground truth annotations (purple). In tissue with sparse nuclei, such as fibroblasts from prostate tissue, NES did not add an additional benefit to DNA alone. In tissues where NES does not exhibit the characteristic nuclear ring, as in glioblastoma, the accuracy was similarly not improved. Scale bar denotes 20 micrometers.

### Combining NES images and real augmentation has a cumulative effect

To determine whether real augmentation and NES would combine during model training to achieve superior segmentation precision relative to the use of either type of data alone, we trained and tested models under four different scenarios (using all three learning frameworks; **Figure 4**). We used images from the small intestine, a tissue containing nuclei having a wide variety of morphologies, and then extended the analysis to other tissue types (see below). Models were evaluated on defocused DNA test data to increase the sensitivity of the experiment. In the first scenario, we trained baseline models using in-focus DNA image data and tested models on unseen in-focus DNA images. With tissues such as the small intestine, which are challenging to segment because they contain densely-packed nuclei, scenario A resulted in slightly under-segmented predictions. In Scenario B and for all subsequent scenarios, defocused DNA images were included in the test set, giving rise to contours that were substantially misaligned with ground truth annotations and resulted in higher undersegmentation. False-positive predictions and imprecise localizations of the nuclei membrane were observed in areas devoid of nuclei and with very low contrast (**Figure 4a**). When NES images were included in the training set (Scenario C), nuclear boundaries were more consistent with ground truth annotations, although false-positive predicted nuclei still remained. The most robust performance across ML frameworks and tissues was observed when NES images and real augmentation were combined: accurate nuclear boundaries were generally well aligned with ground truth annotations in both shape and in size. Observable differences in the placement of segmentation masks were reflected in improvements in IoU: for all three deep learning frameworks, including NES data and real augmentations increased the fraction of nuclei retained by 50% at an IoU threshold of 0.6 (**Figure 4b**). The accuracy of UnMICST-P (blue curve) trained on in-focus DNA data alone was higher than the other two baseline models at all IoU thresholds, suggesting that UnMICST-P has a greater capacity to learn. UnMICST-P may have an advantage in experiments in which staining the nuclear envelope proves difficult or impossible.

**Figure 4:**
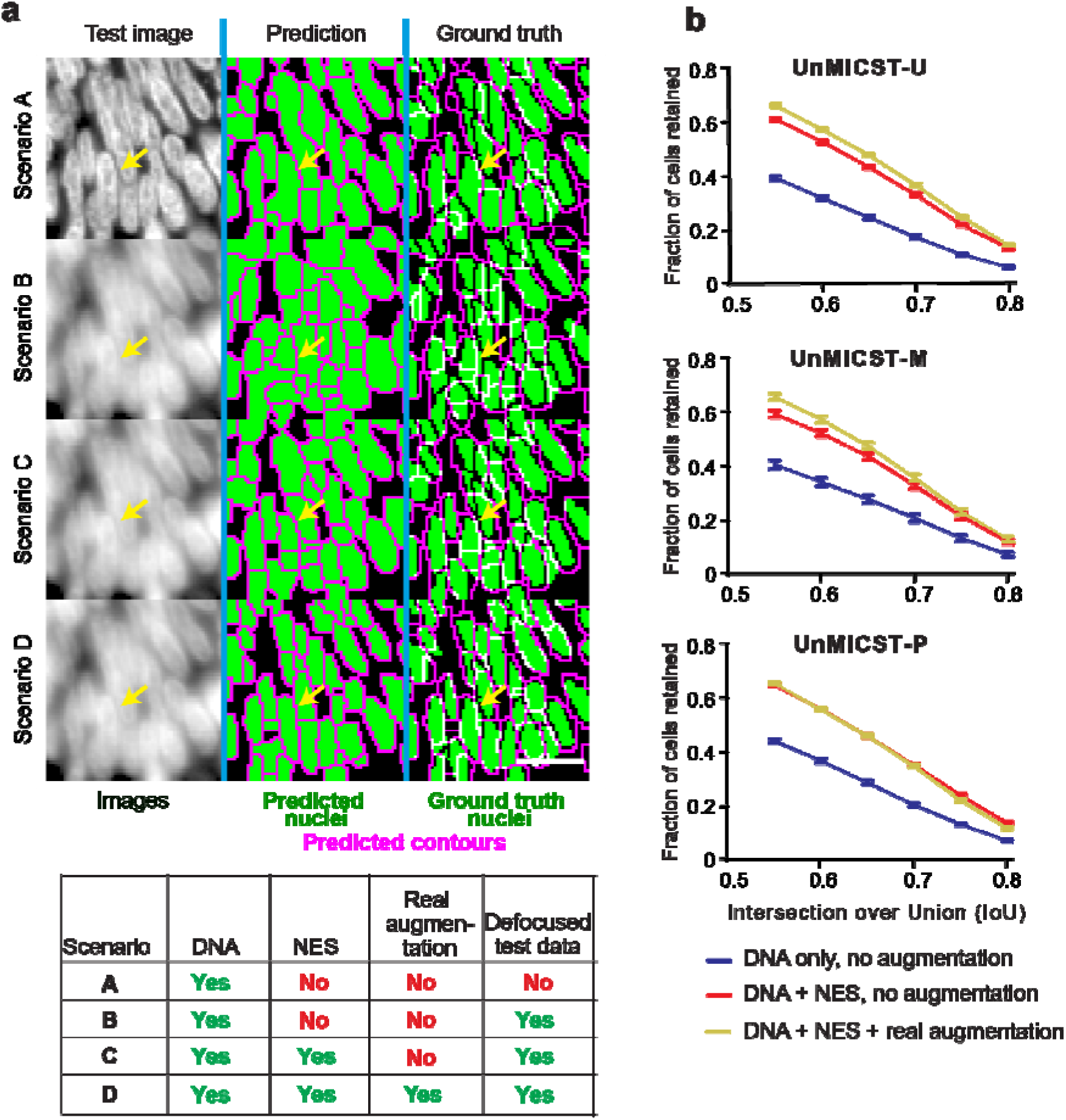
Combination of NES and real image augmentations on segmentation performance. NES - nuclear envelope staining. **a**) Models trained with in-focus DNA data alone produced probability maps that were undersegmented, especially in densely-packed tissue such as small intestine (Scenario A). When tested on defocused data, nuclei borders were largely incorrect (Scenario B). Adding NES restored nuclei border shapes (Scenario C). Combining NES and real augmentations reduced false positive detections and produced nuclei masks better resembling the ground truth labels (Scenario D). Scalebar denotes 20 micrometers. Table legend shows conditions used for each scenario A-D. Yellow arrow indicates a blurry cell of interest where accuracy improves with NES and real augmentation **b**) Graphs compare the accuracy represented as the number of cells retained across varying IoU thresholds with all models from UnMICST-U (top), UnMICST-M (center), and UnMICST-P (bottom). In all models, more nuclei were retained when NES and real augmentations were used together during training (yellow curves) compared to using NES without real augmentations (red curves) or DNA alone (blue curves).

### Combining NES and real augmentation is advantageous across multiple tissue types

To determine if improvements in segmentation would extend to multiple tissue types we repeated the analysis described above using three scenarios for training and testing with both in-focus (**Figure 5a**) and defocused images (**Figure 5b**). Scenario 1 used in-focus DNA images for training (blue bars), scenario 2 used in-focus DNA and NES images (red bars), and scenario 3 used in-focus DNA and NES images plus real augmentation (green bars). While the magnitude of the improvement varied with tissue type and test set (panel a vs b), the results as a whole support the conclusion that including both NES and real augmentations during model training confers statistically significant improvement in segmentation accuracy with multiple tissue types and models. The accuracy boost was greatest when models performed poorly (e.g., in scenario 1 where models were tested on defocused colon image data; **Figure 5b**, blue bars), so that segmentation accuracy became relatively uniform across tissue and cell types. As a final test, we re-examined the whole slide melanoma image described above (which had not been included in any training data) and evaluated IoU, AP and F1-scores. The data were consistent regardless of metric and showed that all three models benefitted from the inclusion of training data that included NES images and real augmentations (**Supplementary Figure 3**). The improvement in accuracy, however, was modest and similar to lung adenocarcinoma. We attribute this to the fact that, like lung adenocarcinoma, melanoma has less dense regions, which our baseline models already performed well on.

**Figure 5:**
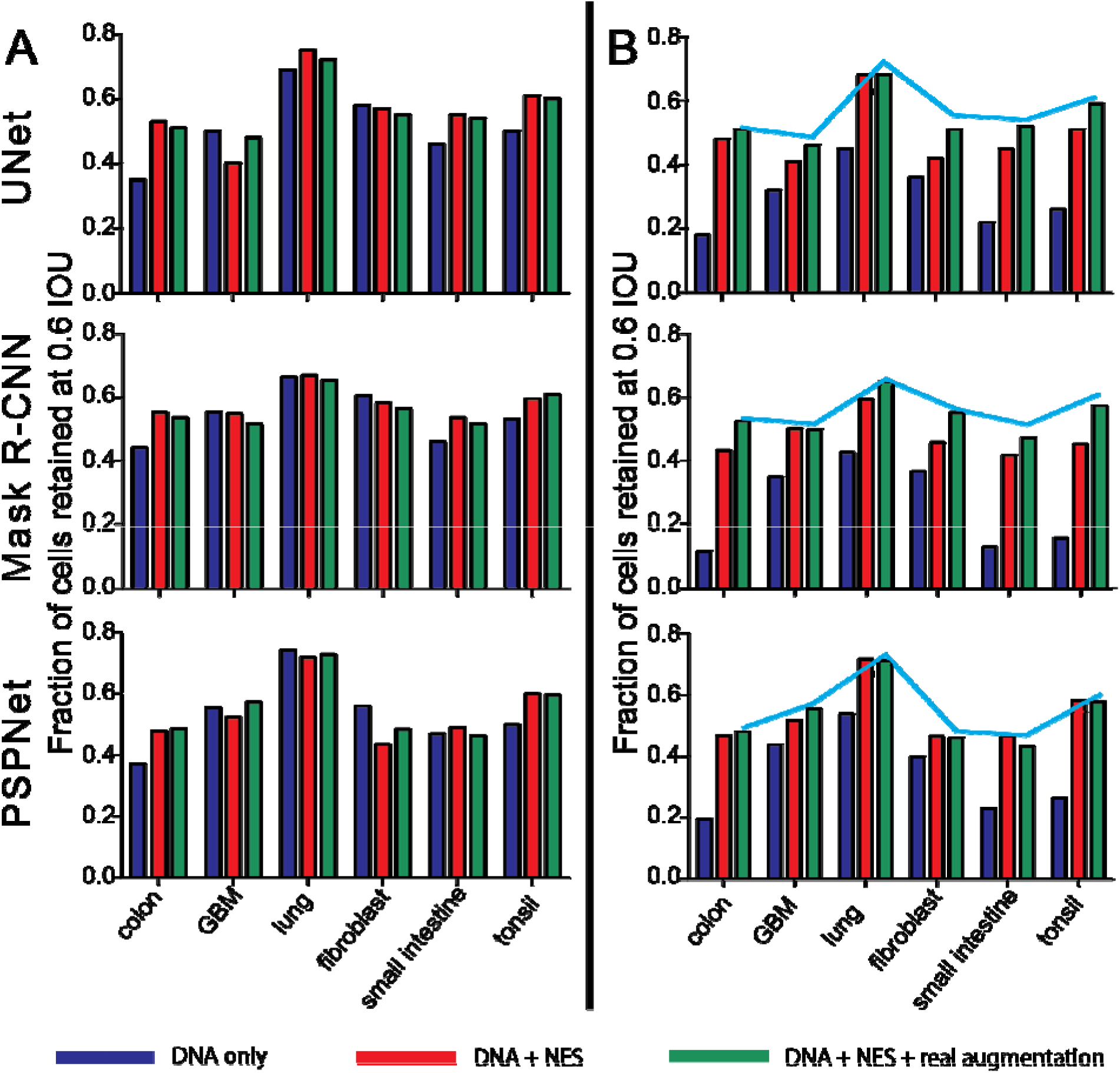
Assessing different training strategies on a) in-focus and b) defocused test data for different tissue types. **a**) In all tissue types apart from GBM, the addition of NES (red bars) and the use of real augmentations combined with NES (green bars) in training data offered superior accuracy compared to using DNA alone (blue bars). **b**) When the models were tested on defocused data, all tissues (including GBM unexpectedly) showed benefits resulting from using NES (red bars) combined with real augmentations (green bars). The line plot indicates highest accuracy achieved for each tissue when tested on in-focus data from panel a.

### Applying UnMICST to highly multiplex whole-slide tissue images

To investigate the overall improvement achievable with a representative UnMICST model, we tested UnMICST-U with and without real or computed augmentations and NES data on all six tissues as a set, including in-focus, saturated and out-focus images (balancing the total amount of training data in each case). A 1.7-fold improvement in accuracy was observed at an IoU of 0.6 for the fully trained model (i.e. with NES data and real augmentations; **Figure 6a**). Inspection of segmentation masks also demonstrated more accurate contours for nuclei across a wide range of shapes. The overall improvement in accuracy was substantially greater than any difference observed between semantic and instance segmentation frameworks. We, therefore, focused subsequent work on the most widely used framework: U-Net.

**Figure 6:**
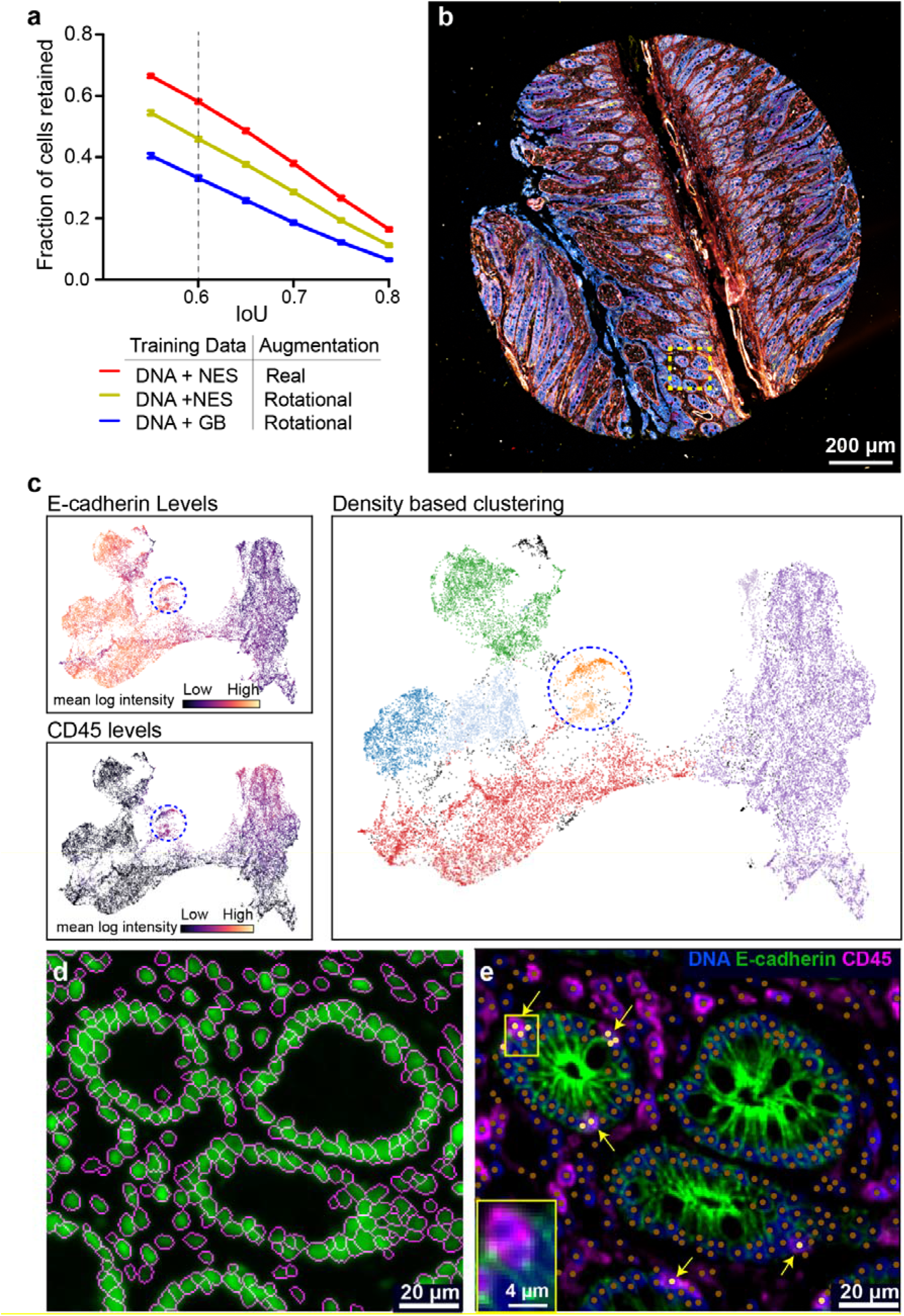
Applying UnMICST models to highly multiplexed image data. **a)** Accuracy improvement of UnMICST-U models trained with and without NES (nuclear envelope staining) as compared to DNA alone, and real augmentations as compared to computed blur (GB; Gaussian blur). To balance training dataset size, GB was substituted for NES data and computed 90/180 degree rotations were substituted for real augmentations. **b)** A 64-plex CyCIF image of a non-neoplastic small intestine TMA core from the EMIT dataset. Dashed box indicates region of interest for panels **d** and **e**. **c)** UMAP projection using single cell staining intensities for 14 marker proteins (see methods). The color of the data points represents the intensity of E-cadherin (top left) or CD45 (bottom left) across all segmented nuclei. Density-based clustering using HDBSCAN identified distinct clusters (each denoted by a different color) that were positive for either E-cadherin or CD45 as well as a small number of double-positive cells (blue dashed circle). **d)** Enlarged region of yellow dashed box from **b** showing segmentation mask outlines (magenta) overlayed onto DNA channel (green). **e)** Composite image of DNA, E-cadherin, and CD45 of the same region. Nuclei centroids from segmentation denoted by brown dots. Cells positive for both E-cadherin and CD45 (from blue dashed circle in panel **c** are marked with yellow arrows and yellow dots. Inset: enlarged view of boxed region showing overlapping immune and epithelial cells.

We also tested a fully trained UnMICST-U model on a 64-plex CyCIF image of non-neoplastic small intestine tissue from the EMIT TMA (**Figure 6b**). Staining intensities were quantified on a per-cell basis, and the results visualized using Uniform Manifold Approximation and Projection (UMAP; **Figure 6c**). Segmentation masks were found to be well-located with little evidence of under or over-segmentation (**Figure 6d**). Moreover, whereas 21% of cells with segmented nuclei stained positive (as determined by using a Gaussian-mixture model) for the immune cell marker CD45, and 53% stained positive for the epithelial cell marker E-cadherin, less than 3% were positive for both. No known cell type is actually positive for both CD45 and E-cadherin, and the very low abundance of these double-positive “cells” is evidence of accurate segmentation. When we examined some of the 830 double positive cells (blue dashed circle in **Figure 6c**) we found multiple examples of a CD3^+^ T cell (yellow arrowheads; light yellow dots in **Figure 6e**) tightly associated with or between the epithelial cells of intestinal villi (green “kiwi” structure visible in **Figure 6e**). This is consistent with the known role of the intestinal epithelium in immune homeostasis^38^. In these cases, the ability of humans to distinguish immune and epithelial cells relies on prior knowledge, multi-dimensional intensity features and subtle differences in shape and texture – none of which were aspects of model training. Thus, future improvements in tissue segmentation are likely to require the development of CNNs able to classify rare but biologically significant spatial arrangements, rather than simple extensions of the general purpose segmentation algorithms described here.

### Some tissues still pose a challenge for nuclei segmentation

Of all the tissue types annotated and tested in this paper, non-neoplastic ovary was the most difficult to segment (**Supplementary Figure 4a**) and addition of ovarian training data to models trained on data from other tissues decreased overall accuracy (**Supplementary Figure 4b**). We have previously imaged ovarian cancers at even higher resolution (60x/1.42NA sampled at 108 nm pixel size); (Färkkilä et al., 2020) using optical sectioning and deconvolution microscopy; inspection of these images reveals nuclei with highly irregular morphology, poor image contrast, and dense packing (**Supplementary Figure 4c**) unlike colon adenocarcinoma (**Supplementary Figure 4d**). Thus, additional research, possibly involving different NES antibodies, will be required to improve performance with ovarian and other difficult to segment tissues. Until then, caution is warranted when combining training data from tissues with very different nuclear morphologies.

## DISCUSSION

This paper makes four primary contributions to the growing literature on the segmentation of tissue images, which is an essential step in single-cell data analysis. First, we consciously included in training and test data the types of focus and intensity artefacts that are commonly encountered in whole-slide images, particularly images of human tissues acquired in the course of clinical care and treatment. This contrasts with other recent papers that focus on optimal fields of view. Second, we show that addition of real augmentations comprising defocused and saturated images to model training data improves segmentation accuracy to a significant extent whereas augmentations based on Gaussian blurring provide minimal benefit. These results extend to deep learning frameworks based on instance segmentation (UnMICST-M) and on semantic segmentation (UnMICST-U and UnMICST-P). Third, we show that it is often possible to increase segmentation accuracy by including additional data (NES) on nuclear envelop morphology, although identifying suitable antibody cocktails is not trivial. Finally, using newly generated labeled training data for multiple tissue types, we show that real augmentation and NES combine to improve the robustness and accuracy of segmentation across many tissues; these improvements are directly applicable to the real-world task of segmenting high dimensional tissue and tumor images. Moreover, the magnitude of improvement observed with NES data or real augmentation is substantially greater than the differences observed between ML frameworks. UnMICST models, therefore, represent a good starting point for performing image segmentation on rapidly growing tissue data repositories. Errors remaining when multiplexed images are segmented using optimized UnMICST models appear to have a subtle biological basis. Development of additional “physiology aware” machine-learning models may be necessary to reduce these apparent errors.

One of the surprises in the current work was the seemingly low level of agreement achieved by two human experts annotating the same image data; we estimated that only 60% of the annotated nuclei between annotators had an overlap of 60% or greater (0.6 IoU threshold). Poor agreement is almost certainly a consequence of our use of a stringent sweeping IoU scoring criterion that measures the fraction of pixels that overlap between two segmentation masks. The alternative, and widely-used F1 score, which determines whether two observers (or an observer and a machine) agree on the of nuclear features, achieves inter-observer and automated segmentation accuracy of 0.78, which is comparable to the highest F1-scoring tissue reported for Mesmer^39^, another deep learning model applied to tissue images. Moreover, our results with IoU values are similar to those recently reported by Kromp et al.^17^ (when IoU tresholds are adjusted to enable direct comparison). The authors of Cellseg^40^ also report comparable segmentation accuracies and note the difficulty of achieving a high IoU value with cells that vary dramatically in shape and focus.

It would therefore appear that many studies have achieved similar levels of inter-observer agreement and that our results are not an outlier, even though we include problematic data. This points to a fundamental challenge for all supervised learning approaches whose solution is not immediately clear. Collection of precise 3D data followed by imposition of different levels of blurring and addition of intensity artefacts will be needed to understand the origins of inter-observer disagreement in tissue images and achieve higher quality training and test data. Despite this concern, it seems likely that practical improvements in segmentation are likely to come from combining recently described advances. For example, Greenwald et al^41^ use a clever community-based approach to acquire much more training data than in the current work, Kromp et al.^17^ combine tissue images with ground truth annotation acquired from cultured cells (by a team of undergraduate students), whereas the current work focuses on the use of NES and real augmentations to improve the robustness of segmentation algorithms across the board.

From a machine learning perspective, the value of adding images to training data is self-evident. Experimental feasibility is not always so clear. A key tradeoff is that the greater the number of fluorescence channels used for segmentation, the fewer the channels available for the collection of data on other markers. Fortunately, the development of highly multiplexed imaging has made this less relevant because collection of 20-40 or more image channels (each corresponding to a different fluorescent antibody) has become routine. This makes it straightforward to reserve two channels for segmentation. The cost-benefit ratio of adding extra segmentation data will be different in high content screening of cells in multi-well plates, for which inexpensive reagents are generally essential than in tissue imaging. In tissues, the morphology of nuclear lamin changes with disease state^42^, cell type, activation state and numerous other biological processes. While this challenges segmentation routines, imaging lamins is also likely to provide valuable biological information, further arguing for routine collection of these data^43^. To allow others to build on the current work, we are releasing all training and test images, their segmentation masks and annotations, and real augmentations for multiple types of tissue (tonsil, ovary, small intestine and cancers of the colon, brain, lung, prostate) via the EMIT resource; models are released as components of the UnMICST model resource (see data and code availability information).

The most immediately generalizable finding from this work is that real augmentation outperforms computed augmentation generated using Gaussian kernels. Blurring and image saturation are an inevitable consequence of the limited bandwidth of optical systems, the thickness of specimens relative to the depth of field, light scattering, diffraction, the use of non-immersion objective lenses and consequent refractive index mismatches, and a variety of other physical processes. Real out-of-focus blur also differs when the focal plane is above and below the specimen. Areas for future application of real augmentations could include inhomogeneous light sources and stage jitter. It will undoubtedly be useful to determine kernels for more effective computed augmentation, but collecting real augmentation data imposes a minimal burden in a real-world setting. Our observation that real augmentation outperforms computed augmentation may also have general significance outside of the field of microscopy: with any high-performance camera system, real out-of-focus data will inevitably be more complicated than Gaussian blur.

## CODE AND DATA AVAILABILITY

To allow others to build on the current work, we are releasing all training, validation and test images, their annotations, and real augmentations for multiple types of tissue (tonsil, ovary, small intestine and cancers of the colon, brain, lung, prostate) via the EMIT resource; models and scripts for training and inference are released as components of the UnMICST model resource (see data and code availability information). https://labsyspharm.github.io/UnMICST-info/

## METHODS

### Sample preparation for imaging

To generate images for model training and testing, human tissue specimens from multiple patients were used to construct a multi-tissue microarray (HTMA427) under an excess (discarded) tissue protocol approved by the Institutional Review Board (IRB) at Brigham and Women’s Hospital (BWH IRB 2018P001627). One or two 1.5 mm diameter cores were taken from tissue regions with the goal of acquiring one or two examples of different healthy or tumor types including non-neoplastic medical diseases and secondary lymphoid tissues such as tonsil. Slides were stained with the following reagents from Cell Signaling Technologies (Beverly MA, USA) and Abcam (Cambridge UK).

Before imaging, slides were mounted with 90% glycerol and a #1.5 coverslip. Prior to algorithmic evaluation, the images were split into three mutually disjoint subsets and used for training, validation, and testing.

### Acquisition of image data and real augmentations

The stained TMA was imaged on a INCell 6000 (General Electric Life Sciences) microscope equipped with a 20x/0.75 objective lens (370 nm nominal lateral resolution at 550 nm wavelength) and a pixel size of 0.325 μm per pixel. Hoechst and lamin-A647 were excited with a 405 nm and 642 nm laser, respectively. Emission was collected with the “DAPI” (455/50 nm) and “Cy5” (682/60 nm) filter sets with exposure times of 60 ms and 100 ms, respectively. Whole-slide imaging involved acquisition of 1,215 tiles with an 8% overlap, which is recommended for stitching in ASHLAR, a next generation stitching and registration algorithm for large images (https://github.com/labsyspharm/ashlar). To generate defocused data, we acquired images from above and below the focal plane by varying the Z-axis by 3 μm in both directions. To generate saturated images of DNA staining, a 150ms exposure time was used. These two types of “suboptimal” data were then used for “real augmentation” during model training, as described below.

Representative cores for lung adenocarcinoma, non-neoplastic small intestine, normal prostate, colon adenocarcinoma, glioblastoma, non-neoplastic ovary, and tonsil were extracted from image mosaics and down-sampled by a factor of 2 to match the pixel size of images routinely acquired and analyzed in MCMICRO^33^. Images were then cropped to 256 x 256-pixel tiles, and in-focus DNA and NES were imported into Adobe Photoshop to facilitate human annotation of nuclear boundaries. Annotations for contours and background classes were labelled on separate layers while swapping between DNA and NES as necessary. To save time, we drew complete contours of nuclei and filled these in using the Matlab *imfill* operation to generate nuclei centers. For nuclei at the image borders where contours would be incomplete, we manually annotated nuclei centers. As described by Ronneberger et al. (2015), a fourth layer was used to mark areas between clumped cells. These additional annotations made it possible to specifically penalize models that incorrectly classified these pixels. Annotations on a held-out test dataset were validated by a second annotator and measured using the F1-score. The F1-score evaluation between both annotated ground truths was high and demonstrated excellent agreement (**Supplementary Figure 1**).

Because original, defocused, and saturated images of DNA were all acquired in the same image stack, it was possible to use a single registered set of DNA annotations across all augmented image channels. To produce the training set, each image was cropped into 64 x 64 patches, normalized to use the full dynamic range, and further augmented using 90-degree rotations, reflections, and 20% upscaling. Consistent with the training set, the validation and test sets also include defocused and saturated examples but were not augmented with standard transformations. The ratio of data examples present in the training, validation, and test set split was 0.36:0.24:0.4. For a fair comparison across models, the same dataset and split were used for the three deep learning frameworks described in this manuscript (**Supplementary Table 2**).

### Model implementation

To facilitate model training, three distinct state-of-the-art architectures were separately, trained, implemented and evaluated. They are, in no particular order, UNet, Mask R-CNN, and PSPNet and were adopted from their original references without modification to their architecture. UNet was selected for its prior success in the biomedical domain, Mask R-CNN was selected for its ability to perform both object detection and mask generation, and PSPNet was selected for its capacity to integrate image features from multiple spatial scales. Training, validation, and test data were derived from 12 cores in 7 tissues and a total of 10,359 nuclei in the composition of colon – 1,142; glioblastoma (GBM) – 675; lung – 1735; ovarian – 956; fibroblast – 922; small intestine – 1677; tonsil – 3252. To maintain consistency of evaluation across segmentation algorithms, segmentation accuracy was calculated by counting the fraction of cells in a held out test set that passed a sweeping Intersection over Union (IoU) threshold. The NES channel was concatenated to the DNA channel as a three-dimensional array as input into each architecture.

### Model Training

#### UnMICST-U models

A three-class UNet model^14^ was trained based on annotation of nuclei centers, nuclei contours, and background. The neural network is comprised of 4 layers and 80 input features. Training was performed using a batch size of 32 with the Adam Optimizer and a learning rate of 0.00005 with a decay rate of 0.98 every 5,000 steps until there was no improvement in accuracy or ~100 epochs had been reached. Batch normalization was used to improve training speed. During training, the bottom layer had a dropout rate of 0.35, and L1 regularization was implemented to minimize overfitting^44, 45^ and early stopping. Training was performed on workstations equipped with NVidia GTX 1080 or NVidia TitanX GPUs.

#### UnMICST-M models

Many segmentation models are based on the Mask R-CNN architecture^15^, Mask R-CNN has previously exhibited excellent performance on a variety of segmentation tasks. Mask R-CNN begins by detecting bounding boxes of nuclei and subsequently performs segmentation within each box. This approach eliminates the need for an intermediate watershed, or equivalent, segmentation step. Thus, Mask R-CNN directly calculates a segmentation mask, significantly reducing the overhead in traditional segmentation pipelines. We adopted a ResNet50^46^ backbone model in the UnMICST-M implementation and initialized the weights using pretrained values from the COCO object instance segmentation challenge^30^ to improve convergence properties. For efficient training, we upsampled the original input images to 800 x 800-pixels and trained a model for 24 epochs using a batch size of 8. The Adam optimizer, with a weight decay of 0.0001 to prevent overfitting, was exploited with a variable learning rate, initially set to 0.01 and decreased by a factor of 0.1 at epochs 16 and 22. Training was performed on a compute node cluster using 4 NVidia TitanX or NVidia Tesla V100 GPUs. For evaluation and comparison, we used the model with the highest performance on the validation set, following standard practice.

#### UnMICST-P models

We trained a three class PSPNet model^47^ to extract cell nuclei centers, nuclei contours, and background from a wide variety of tissue types. PSPNet is one of the most widely used convolutional neural networks for the semantic segmentation of natural scene images in the computer vision field. The network employs a so-called pyramid pooling module whose purpose is to learn global as well as local features. The additional contextual information used by PSPNet allowed the segmentation algorithm to produce realistic probability maps with greater confidence. We used ResNet101 as a backbone. Training of the network was performed using a batch size of 8 with an image size of 256 x 256-pixels for 15,000 iterations or until the minimum loss model was obtained. A standard cross entropy loss function was used during training. Gradient descent was performed using the Adam optimizer with a learning rate of 0.0001 and a weight decay parameter of 0.005 via L2 regularization. Batch normalization was employed for faster convergence, and a dropout probability of 0.5 was used in the final network layer to mitigate overfitting. The model training was performed on a compute cluster node equipped with NVidia Tesla V100 GPUs.

### Analysis of multi-dimensional data

For the analysis shown in Figure 6, a 64-plex CyCIF image of non-neoplastic small intestine tissue from the EMIT TMA (https://www.synapse.org/#!Synapse:syn22345748/) was stained with a total of 45 antibodies as described in protocols https://www.protocols.io/view/ffpe-tissue-pre-treatment-before-t-cycif-on-leica-bji2kkge and https://dx.doi.org/10.17504/protocols.io.bjiukkew. Images were segmented using the UnMICST-U model trained on DNA with NES data and real augmentations. Mean fluorescence intensities across 45 markers for 27,847 segmented nuclei were quantified as described in^33^. E-cadherin positive and CD45 positive cells were identified using Gaussian-mixture models on log-transformed data. For multivariate clustering, log-transformed mean intensities of all single cells of 14 selected protein markers (E-cadherin, pan-cytokeratin, CD45 CD4, CD3D, CD8, RF3, PML, GLUT1, GAPDH TDP43, OGT, COLL4, an EPCAM) were pre-processed using Uniform Manifold Approximation and Projection (UMAP)^48^ and clustered using Hierarchical Density-Based Spatial Clustering of Applications with Noise (HDBSCAN)^49^. Clusters expressing a high level of both E-cadherin and CD45 were identified and overlaid on a false-colored image showing the staining of DNA, E-cadherin, and CD45.

## ACKNOWLEDGEMENTS

We thank Alyce Chen and Madison Tyler for their help with this manuscript. The work was funded by NIH grants U54-CA225088 and U2C-CA233262 to P.K.S. and S.S and by the Ludwig Cancer Center at Harvard. Z.M. is supported by NCI grant R50-CA252138. We thank Dana-Farber/Harvard Cancer Center (P30-CA06516) for the use of its Specialized Histopathology Core.

## AUTHOR CONTRIBUTIONS

The study design was conceived by CY, WDJ, EN, and PKS. Image acquisition and annotation was done by CY. SS provided the EMIT TMA sample and validated the tissue types. TMA staining was performed by ZM and CAJ. Data analysis was performed by CY, WDJ, EN, and YAC. YAC and CY performed the single cell quantitative analysis and analysis found in figure 6. Additional coding was done by MC. Additional experiments were conducted by DW. PKS, SS, HP supervised the study. All authors contributed to the writing and editing of the manuscript.

## OUTSIDE INTERESTS

PKS is a member of the SAB or BOD member of Applied Biomath, RareCyte Inc., and Glencoe Software, which distributes a commercial version of the OMERO database; PKS is also a member of the NanoString SAB. In the last five years the Sorger lab has received research funding from Novartis and Merck. Sorger declares that none of these relationships have influenced the content of this manuscript. SS is a consultant for RareCyte Inc. The other authors declare no outside interests.

## Notes

### Summary of Updates

-Additional segmentation metrics added -Additional author added -Interobserver agreement metric added -Supplementary file updated

## REFERENCES

1. Gerdes, M. J. et al. Highly multiplexed single-cell analysis of formalin-fixed, paraffin-embedded cancer tissue. Proc. Natl. Acad. Sci. U. S. A. 110, 11982–11987 (2013).

2. Giesen, C. et al. Highly multiplexed imaging of tumor tissues with subcellular resolution by mass cytometry. Nat. Methods 11, 417–422 (2014).

3. Angelo, M. et al. Multiplexed ion beam imaging of human breast tumors. Nat. Med. 20, 436–442 (2014).

4. Lin, J.-R. et al. Highly multiplexed immunofluorescence imaging of human tissues and tumors using t-CyCIF and conventional optical microscopes. eLife 7, e31657 (2018).

5. Stack, E. C., Wang, C., Roman, K. A. & Hoyt, C. C. Multiplexed immunohistochemistry, imaging, and quantitation: A review, with an assessment of Tyramide signal amplification, multispectral imaging and multiplex analysis. Methods 70, 46–58 (2014).

6. Immunologists, A. A. of. The Demonstration of Pneumococcal Antigen in Tissues by the Use of Fluorescent Antibody. J. Immunol. 45, 159–170 (1942).

7. Albertson, D. G. Gene amplification in cancer. Trends Genet. 22, 447–455 (2006).

8. Shlien, A. & Malkin, D. Copy number variations and cancer. Genome Med. 1, 62 (2009).

9. Amin, M. B. et al. The Eighth Edition AJCC Cancer Staging Manual: Continuing to build a bridge from a population-based to a more ‘personalized’ approach to cancer staging. CA. Cancer J. Clin. 67, 93–99 (2017).

10. Achim, K. et al. High-throughput spatial mapping of single-cell RNA-seq data to tissue of origin. Nat. Biotechnol. 33, 503–509 (2015).

11. Slyper, M. et al. A single-cell and single-nucleus RNA-Seq toolbox for fresh and frozen human tumors. Nat. Med. 26, 792–802 (2020).

12. McQuin, C. et al. CellProfiler 3.0: Next-generation image processing for biology. PLOS Biol. 16, e2005970 (2018).

13. LeCun, Y., Bengio, Y. & Hinton, G. Deep learning. Nature 521, 436–444 (2015).

14. Ronneberger, O., Fischer, P. & Brox, T. U-Net: Convolutional Networks for Biomedical Image Segmentation. ArXiv150504597 Cs (2015).

15. He, K., Gkioxari, G., Dollár, P. & Girshick, R. Mask R-CNN. ArXiv170306870 Cs (2018).

16. Caicedo, J. C. et al. Nucleus segmentation across imaging experiments: the 2018 Data Science Bowl. Nat. Methods 16, 1247–1253 (2019).

17. Kromp, F. et al. An annotated fluorescence image dataset for training nuclear segmentation methods. Sci. Data 7, 262 (2020).

18. Schwendy, M., Unger, R. E. & Parekh, S. H. EVICAN—a balanced dataset for algorithm development in cell and nucleus segmentation. Bioinformatics 36, 3863–3870 (2020).

19. Schüffler, P. J. et al. Automatic single cell segmentation on highly multiplexed tissue images. Cytometry A 87, 936–942 (2015).

20. Arganda-Carreras, I. et al. Trainable Weka Segmentation: a machine learning tool for microscopy pixel classification. Bioinformatics 33, 2424–2426 (2017).

21. Berg, S. et al. ilastik: interactive machine learning for (bio)image analysis. Nat. Methods 16, 1226–1232 (2019).

22. Aeffner, F. et al. Introduction to Digital Image Analysis in Whole-slide Imaging: A White Paper from the Digital Pathology Association. J. Pathol. Inform. 10, 9 (2019).

23. Lin, J.-R. et al. Multiplexed 3D atlas of state transitions and immune interactions in colorectal cancer. bioRxiv 2021.03.31.437984 (2021) doi:10.1101/2021.03.31.437984.

24. Krizhevsky, A., Sutskever, I. & Hinton, G. E. ImageNet Classification with Deep Convolutional Neural Networks. in Advances in Neural Information Processing Systems 25 (eds. Pereira, F., Burges, C. J. C., Bottou, L. & Weinberger, K. Q.) 1097–1105 (Curran Associates, Inc., 2012).

25. Ahmed Raza, S. E. et al. MIMO-Net: A multi-input multi-output convolutional neural network for cell segmentation in fluorescence microscopy images. in 2017 IEEE 14th International Symposium on Biomedical Imaging (ISBI 2017) 337–340 (IEEE, 2017). doi:10.1109/ISBI.2017.7950532.

26. Shorten, C. & Khoshgoftaar, T. M. A survey on Image Data Augmentation for Deep Learning. J. Big Data 6, 60 (2019).

27. Horwath, J. P., Zakharov, D. N., Mégret, R. & Stach, E. A. Understanding important features of deep learning models for segmentation of high-resolution transmission electron microscopy images. Npj Comput. Mater. 6, 1–9 (2020).

28. Gurari, D. et al. How to Collect Segmentations for Biomedical Images? A Benchmark Evaluating the Performance of Experts, Crowdsourced Non-experts, and Algorithms. 2015 IEEE Winter Conf. Appl. Comput. Vis. (2015) doi:10.1109/WACV.2015.160.

29. Deng, J. et al. ImageNet: A large-scale hierarchical image database. in 2009 IEEE Conference on Computer Vision and Pattern Recognition 248–255 (2009). doi:10.1109/CVPR.2009.5206848.

30. Lin, T.-Y. et al. Microsoft COCO: Common Objects in Context. in Computer Vision – ECCV 2014 (eds. Fleet, D., Pajdla, T., Schiele, B. & Tuytelaars, T.) 740–755 (Springer International Publishing, 2014). doi:10.1007/978-3-319-10602-1_48.

31. Skinner, B. M. & Johnson, E. E. P. Nuclear morphologies: their diversity and functional relevance. Chromosoma 126, 195–212 (2017).

32. Dalle, J.-R. et al. Nuclear pleomorphism scoring by selective cell nuclei detection. in IEEE Workshop on Applications of Computer Vision (WACV 2009), 7-8 December, 2009, Snowbird, UT, USA (IEEE Computer Society, 2009).

33. Schapiro, D. et al. MCMICRO: a scalable, modular image-processing pipeline for multiplexed tissue imaging. Nat. Methods 1–5 (2021) doi:10.1038/s41592-021-01308-y.

34. Nirmal, A. J. et al. The spatial landscape of progression and immunoediting in primary melanoma at single cell resolution. bioRxiv 2021.05.23.445310 (2021) doi:10.1101/2021.05.23.445310.

35. Fischer, E. G. Nuclear Morphology and the Biology of Cancer Cells. Acta Cytol. 64, 511–519 (2020).

36. Kros, J. M. Grading of Gliomas: The Road From Eminence to Evidence. J. Neuropathol. Exp. Neurol. 70, 101–109 (2011).

37. Louis, D., Ohgaki, H., Wiestler, O. & Cavenee, W. WHO Classification of Tumours of the Central Nervous System. (2016).

38. Allaire, J. M. et al. The Intestinal Epithelium: Central Coordinator of Mucosal Immunity. Trends Immunol. 39, 677–696 (2018).

39. Greenwald, N. F. et al. Whole-cell segmentation of tissue images with human-level performance using large-scale data annotation and deep learning. http://biorxiv.org/lookup/doi/10.1101/2021.03.01.431313 (2021) doi:10.1101/2021.03.01.431313.

40. Lee, M. Y. et al. CellSeg: a robust, pre-trained nucleus segmentation and pixel quantification software for highly multiplexed fluorescence images. BMC Bioinformatics 23, 46 (2022).

41. Greenwald, N. F. et al. Whole-cell segmentation of tissue images with human-level performance using large-scale data annotation and deep learning. bioRxiv 2021.03.01.431313 (2021) doi:10.1101/2021.03.01.431313.

42. Sakthivel, K. M. & Sehgal, P. A Novel Role of Lamins from Genetic Disease to Cancer Biomarkers. Oncol. Rev. 10, 309 (2016).

43. Bell, E. S. & Lammerding, J. Causes and consequences of nuclear envelope alterations in tumour progression. Eur. J. Cell Biol. 95, 449–464 (2016).

44. Ng, A. Y. Feature selection, L1 vs. L2 regularization, and rotational invariance. in Proceedings of the twenty-first international conference on Machine learning 78 (Association for Computing Machinery, 2004). doi:10.1145/1015330.1015435.

45. Srivastava, N., Hinton, G., Krizhevsky, A., Sutskever, I. & Salakhutdinov, R. Dropout: a simple way to prevent neural networks from overfitting. J. Mach. Learn. Res. 15, 1929–1958 (2014).

46. He, K., Zhang, X., Ren, S. & Sun, J. Deep Residual Learning for Image Recognition. in 2016 IEEE Conference on Computer Vision and Pattern Recognition (CVPR) 770–778 (2016). doi:10.1109/CVPR.2016.90.

47. Zhao, H., Shi, J., Qi, X., Wang, X. & Jia, J. Pyramid Scene Parsing Network. ArXiv161201105 Cs (2017).

48. Becht, E. et al. Dimensionality reduction for visualizing single-cell data using UMAP. Nat. Biotechnol. 37, 38–44 (2019).

49. Campello, R. J. G. B., Moulavi, D. & Sander, J. Density-Based Clustering Based on Hierarchical Density Estimates. in Advances in Knowledge Discovery and Data Mining (eds. Pei, J., Tseng, V. S., Cao, L., Motoda, H. & Xu, G.) 160–172 (Springer, 2013). doi:10.1007/978-3-642-37456-2_14.

