## supplementary figures and tables for "UnMICST: Deep learning with real augmentation for robust segmentation of highly multiplexed images of human tissues"

**SUPPLEMENTARY MATERIALS**

### Supplementary Figure 1

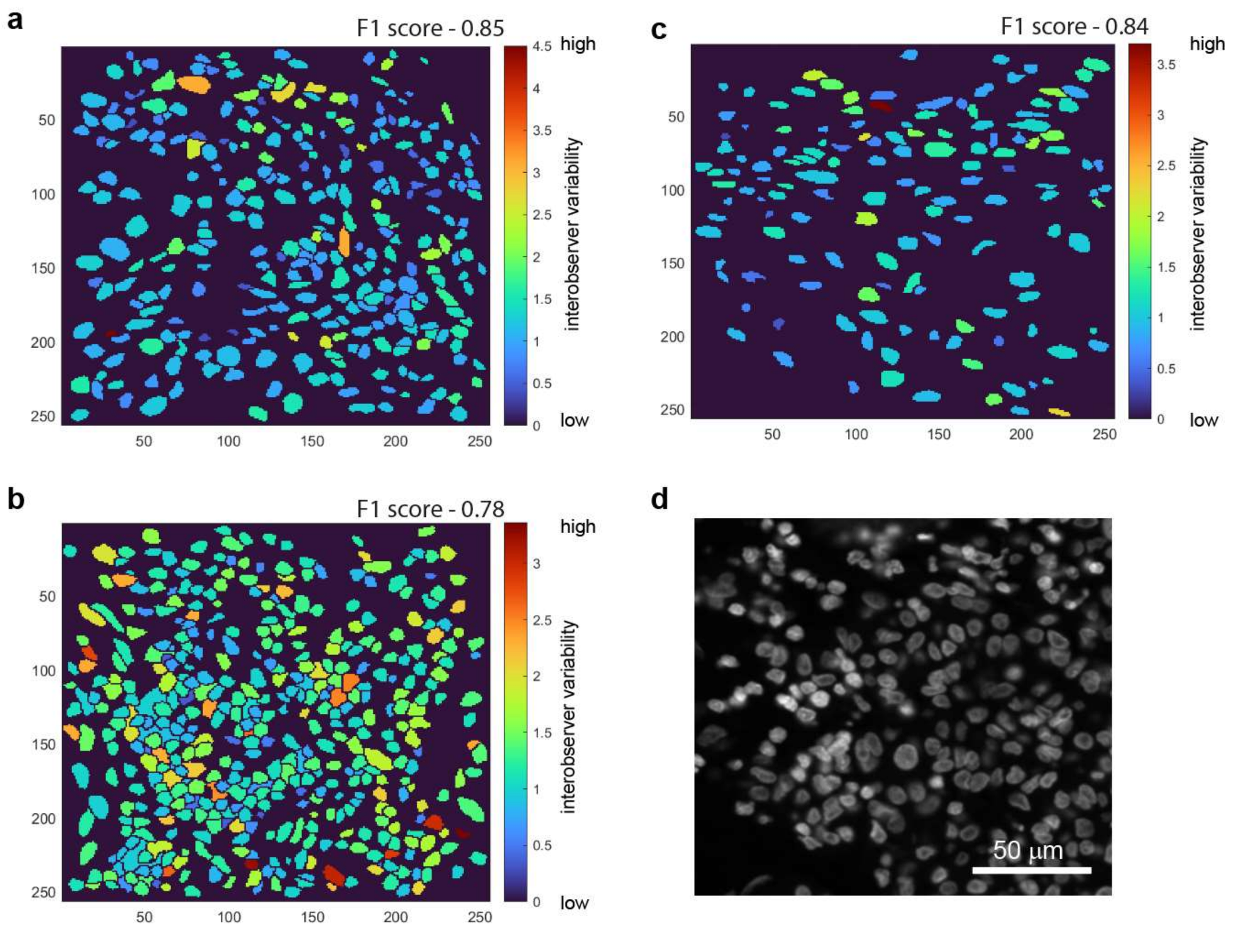

**Supplementary Figure 1: On the comparison of inter-annotator variability.** a-c) Three examples representing differences in manually drawn ground truth segmentation masks between two experienced annotators (who work with microscopy images on a daily basis). An average F1-score of 0.78 between the ground truth masks was obtained, implying that the annotations maintain significant overlap between annotators. The area fold-change per cell drawn across annotators is represented by a color bar, where a red hue implies a larger discrepancy and a blue hue implies a larger agreement. d) Hoechst-stained nuclei image of a) for comparison. Scale bar is 50 μm.

### Supplementary Figure 2

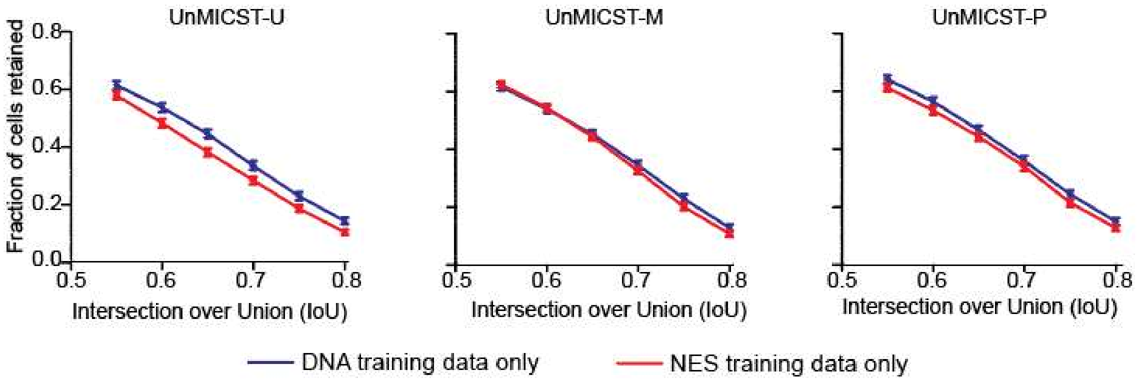

**Supplementary Figure 2: NES data does not substitute for DNA in model training.** Test results when training was performed on DNA (blue curve) and lamin (red curve) individually as single channels for **a**) UnMICST-U, **b**) UnMICST-M, and **c**) UnMICST-P.

### Supplementary Figure 3

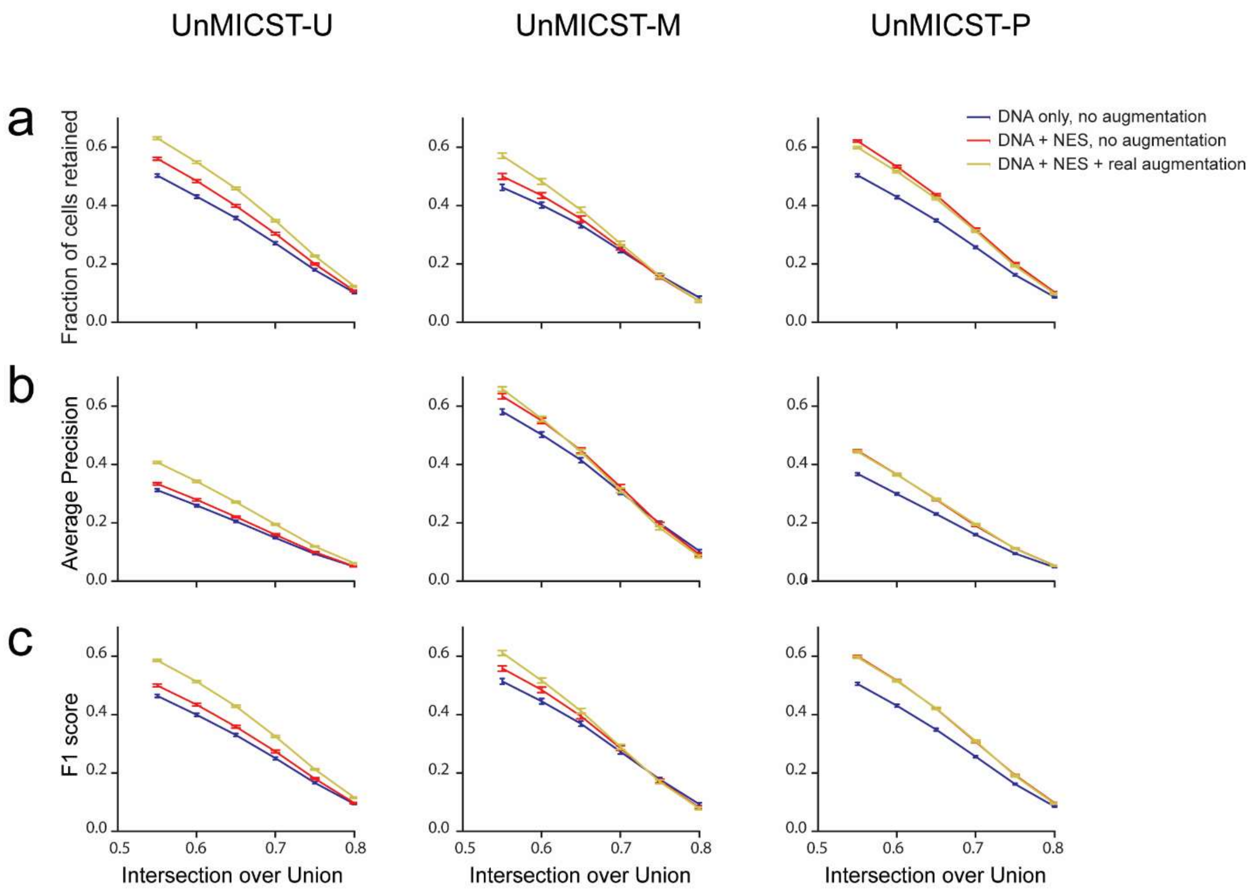

**Supplementary Figure 3: Assessing the use of NES and real augmentations in UnMICST models on unseen test data.** All three learning frameworks in the UnMICST family of models were evaluated on out-of-sample melanoma tissue image data not included in the original training set. NES and real augmentations both have the ability to improve model accuracy on unseen test data in melanoma images. The model performance was assessed using both pixel- and object-level metrics via the (a) fraction of correctly segmented cells, b) Average Precision, and c) F1 score over stringent variable Intersection over Union thresholds. In all three frameworks, the DNA + NES + real augmentations model performance (yellow curves) continues to match or exceed that of the DNA+NES model (red curves) and the DNA-only model (blue curves).

### Supplementary Figure 4

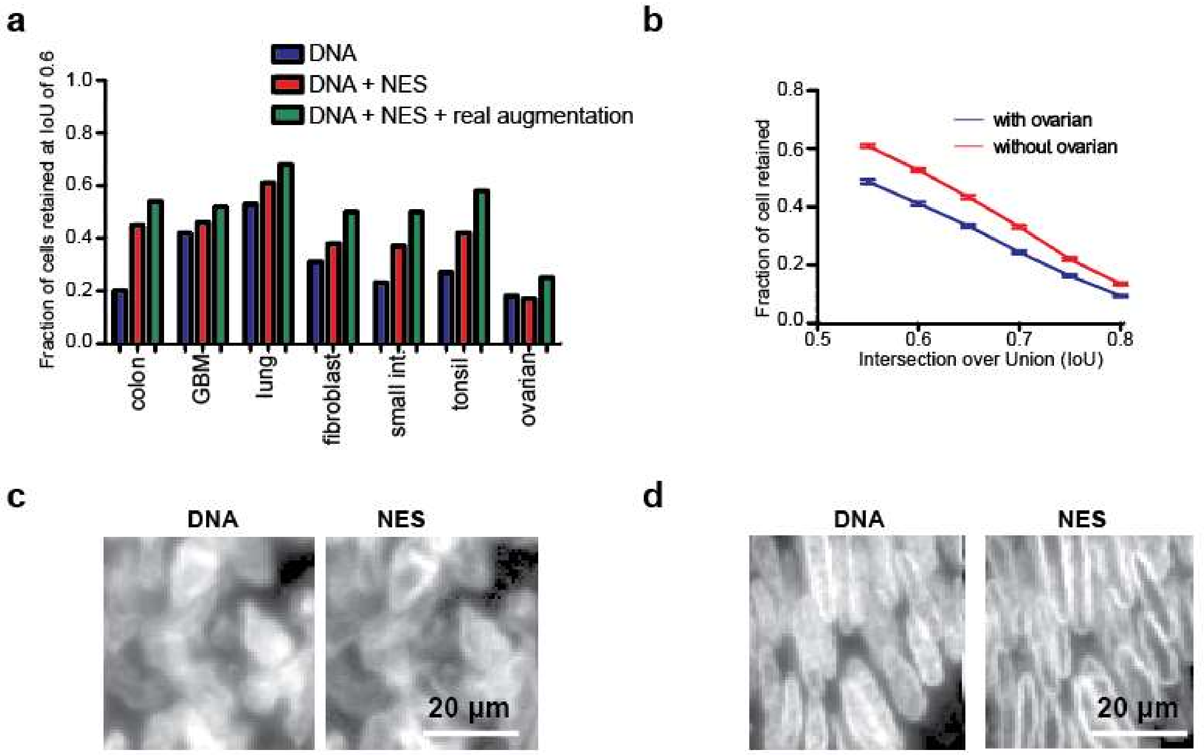

**Supplementary Figure 4: Assessing the use of normal ovarian data in model training**. **a**) Normal ovarian tissue performed the worst with respect to segmentation accuracy among all annotated tissues. Using NES and real augmentations conferred a modest benefit. **b**) The segmentation accuracy across all tissues was lower when ovarian data was included in the training set (blue curve) as opposed to excluded (red curve). **c**) Images showing that annotation of normal ovarian nuclei from DNA is a challenging task, even with NES staining at four times the nominal resolution (0.325 microns per pixel and with a 20x/0.75 objective lens). **d**) In contrast, colon was straightforward to annotate, particularly with NES staining, which forms a distinct halo around the nuclear periphery.

**Supplementary Table 1: High-plex tissue imaging methods**

| | | | |
|---|---|---|---|
| Optical | Fluorophore-based | Single staining cycle but with sequential data acquisition | CO-Detection by indEXing (CODEX)[2]<br><br>Immunostaining with Signal Amplification By Exchange Reaction (Immuno-SABER)[6] |
| | | Cyclic staining and sequential data detection | Cyclic Immunofluorescence (CyCIF)[7]<br><br>Multiplex immunofluorescence (MxIF)[3]<br><br>Iterative indirect immunofluorescence imaging (4i)[9] |
| | Enzyme-based | | Multiplex Immunohistochemistry (mIHC)[4]<br><br>Multiplexed immunohistochemical consecutive staining on single slide (MICSSS)[8] |
| Non-optical | Detection based on atomic mass spectrometry of metal-labelled antibodies | | Imaging Mass Cytometry (IMC)[1]<br><br>Multiplexed Ion Beam Imaging (MIBI)[5] |

**Supplementary Table 2**: Data set size and composition across disjoint training and testing data splits. The model used for training is indicated in the left-hand column and was identical across all three segmentation architectures.

| **Model** | **Training Set Size** | **Test Set Size** |
|---|---|---|
| In-focus DNA | 3636 | 217 |
| In-focus DNA + NES | 3636 | 217 |
| In-focus DNA + Real Augmentations | 21,816 | 1,302 |
| In-focus DNA + NES + Real Augmentations | 21,816 | 1,302 |
| In-focus DNA + Gaussian blur | 14,544 | 868 |
| DNA + Real Augmentations | 5,100 | 645 |
| DNA + 90º/180º rotations | 5,100 | 645 |
| DNA + NES + Real Augmentations | 5,100 | 645 |
| DNA + NES + 90º/180º rotations | 5,100 | 645 |

### Supplementary Note 1: Historical information on segmentation of microscopy images

One of the earliest methods to extract cells from the background involves specifying a global threshold based on the Otsu method[1] in which a threshold is selected that maximizes intraclass variance between two classes. For cell objects that remain clumped and unseparated after simple thresholding, the Otsu method is usually replaced with marker-controlled watershed segmentation. This entails a distance transform operation on the clumped objects to identify a single central 'seed' point per object. The presence of multiple seed points per object (cell) results in oversegmentation and this can usually be resolved with blurring followed by a regional maxima operation. Other methods to identify markers include, but are not limited to, graph cuts or filtering with a blob detector (usually a Laplacian of Gaussian filter)[2–4], level sets followed by mean-shift clustering[5], or a filter bank of rings[6]. Many of these methods have been applied to both fluorescence and H&E images of tissues and while ground truth labeled datasets are not required, fairly extensive parameter tuning and empirical testing is necessary.

A variety of neural network, and other sophisticated, architectures have been explored to improve detection and classification accuracy and expand generalizability to new types of images. Several deep learning architectures developed for natural images have been adapted for marker detection in images of cells including Fully Convolutional Networks (FCNs)[7], Visual Geometry Group (VGG16)[8,9], Residual Networks (ResNets)[10], UNet[11–17], and Mask R-CNN[18,19,19–22]. In classical image analysis, advances in methodology commonly involve the development of new algorithms; any changes in parameter settings needed to accommodate new data are regarded as project-specific details. In contrast, in machine learning approaches, advances in algorithms, learned models and labelled data are all significant; this is particularly true as differentiable architectures for deep learning become increasingly standardized and the quality of trained models is a key point of comparison. While machine learning algorithms outperform classical methods in many image processing applications, model training and evaluation remain substantial challenges in the biomedical domain.

*Biomedical Image datasets*

Image recognition models are commonly trained on diverse sets of natural images, and transfer-style learning is used to improve performance on specific domains. The introduction of the ImageNet dataset[23] catalyzed the application of deep learning to image analysis by providing a set of real-world images along with ground truth annotations. Subsequent applications of ImageNet make clear that machine learning models are only as strong as the underlying training and validation data. In biomedical research, segmentation methods for cell lines[21,24–26] have benefited from large public datasets such as Expert Visual Cell ANnotation (EVICAN)[27], Cellpose[28], the Broad Bioimage Benchmark Collection (BBBC)[29], The Cancer Genome Atlas (TCGA)[30], and Kaggle datasets[31].

Few human-labelled datasets are currently available for highly multiplexed tissue images and existing learned segmentation models generally apply only to specific tissue types[2–6,25,32–35]. Moreover, in most existing datasets, images comprise H&E and not fluorescence images. One exception is a broad 23-tissue study that is part of TCGA, however, these annotations are not publicly available. Moreover, because only the center of the nuclei (as opposed to the nuclear boundary) was marked, TCGA annotations are more suited to cell counting as opposed to segmentation. To address this issue, this current manuscript includes a set of tissue images and densely labelled ground-truth annotations of whole nuclei.

*Image augmentation to improve model training*

Deep learning models have a high capacity to learn large numbers of features. This results in high accuracy but with a substantial danger of overfitting. The use of augmentation methods helps to remedy overfitting by increasing the diversity of the training data[36]. The most common method employs a combination of translational shifts, 90 degree rotations, reflections, and affine transformations[7,15,21,37]. A few studies have also added elastic deformations using B-splines[26,38,39]. These methods are not unique to microscopy images, however, and only a few studies have used augmentation to address variation in the brightness and contrast of otherwise identical images[17] or added synthetically generated camera noise and non-cellular debris to make model training less sensitive to artefacts[13,31]. A particularly interesting form of augmentation used by Kromp *et al.* (2019) involved manually separating cells from the background and arranging nuclei in grids with random positions and orientations, effectively generating new training examples. The authors found that imposing a grid structure did not substantially improve accuracy because cut-out nuclei had hard edges that are atypical of fluorescence images. The authors, therefore, used a generative model trained to relate images of nuclei to their artificial counterparts – nuclei that had been cut out and placed back on their original positions.

To date, common artefacts such as image saturation and defocus have been addressed computationally using image histogram modification and Gaussian blurring[31,40] as well as deliberate defocusing of an actual microscope[29]. The BBBC described by Ljosa et al. is primarily derived from tissue culture cells or transmission and differential interference contrast (DIC) microscopy of model organisms as opposed to human tissues. Ground truth information in Ljosa et al. was also generated automatically as opposed to by human annotation, where the latter is likely to be more accurate.

*Use of stains to aid in segmentation*

In fluorescence microscopy, nuclei are most commonly stained using intercalating dyes such as DAPI or Hoechst 33342[13,19,32,41], SiR-DNA[31], TO-PRO[2], and hematoxylin[3–5,9,10,15,32,34]. When expression of recombinant proteins is feasible (e.g., in cell lines), cells expressing fluorescent protein fusions to histones[42] or spindle components is an effective means to label nuclei[8,14]; this has also been done in genetically engineered mouse models[43], but is not relevant to the analysis of human tissues. In the case of brightfield or phase contrast imaging, nuclear labels are not utilized[27,44,45], but nuclei can often be identified based on brightness. Almost all of these studies have used data from a single imaging channel for nuclei localization, which can be problematic when nuclei are diffuse or close together, both of which are common in cancer specimens. In general, the results of segmentation are superior using fluorescence as opposed to brightfield data[46], but the use of additional channels to more precisely define nuclear boundaries by staining for nuclear lamins or nucleoporins, for example, has not been broadly explored.

### Supplementary References

1. Otsu, N. A Threshold Selection Method from Gray-Level Histograms. (1979) doi:10.1109/TSMC.1979.4310076.
2. Byun, J. *et al.* Automated tool for the detection of cell nuclei in digital microscopic images: application to retinal images. *Mol. Vis.* **12**, 949–960 (2006).
3. Xu, H., Lu, C. & Mandal, M. An efficient technique for nuclei segmentation based on ellipse descriptor analysis and improved seed detection algorithm. *IEEE J. Biomed. Health Inform.* **18**, 1729–1741 (2014).
4. Xu, H., Lu, C., Berendt, R., Jha, N. & Mandal, M. Automatic Nuclei Detection Based on Generalized Laplacian of Gaussian Filters. *IEEE J. Biomed. Health Inform.* **21**, 826–837 (2017).

5. Qi, X., Xing, F., Foran, D. J. & Yang, L. Robust Segmentation of Overlapping Cells in Histopathology Specimens Using Parallel Seed Detection and Repulsive Level Set. *IEEE Trans. Biomed. Eng.* **59**, 754–765 (2012).
6. Liu, C., Shang, F., Ozolek, J. A. & Rohde, G. K. Detecting and segmenting cell nuclei in two-dimensional microscopy images. *J. Pathol. Inform.* **7**, 42 (2016).
7. Lux, F. & Matula, P. Cell Segmentation by Combining Marker-Controlled Watershed and Deep Learning. *ArXiv200401607 Cs Eess* (2020).
8. Wang, C. *et al.* Deep learning pipeline for cell edge segmentation of time-lapse live cell images. *bioRxiv* 191858 (2019) doi:10.1101/191858.
9. Shahzad M *et al.* Robust Method for Semantic Segmentation of Whole-Slide Blood Cell Microscopic Images. *Comput. Math. Methods Med.* **2020**, 4015323–4015323 (2020).
10. Lee, H. & Jeong, W.-K. Scribble2Label: Scribble-Supervised Cell Segmentation via Self-Generating Pseudo-Labels with Consistency. *ArXiv200612890 Cs* (2020).
11. Al-Kofahi, Y., Zaltsman, A., Graves, R., Marshall, W. & Rusu, M. A deep learning-based algorithm for 2-D cell segmentation in microscopy images. *BMC Bioinformatics* **19**, 365 (2018).
12. McQuin, C. *et al.* CellProfiler 3.0: Next-generation image processing for biology. *PLOS Biol.* **16**, e2005970 (2018).
13. Schmidt, U., Weigert, M., Broaddus, C. & Myers, G. Cell Detection with Star-convex Polygons. *ArXiv180603535 Cs* **11071**, 265–273 (2018).
14. Wen, C. *et al.* Deep-learning-based flexible pipeline for segmenting and tracking cells in 3D image time series for whole brain imaging. *bioRxiv* 385567 (2018) doi:10.1101/385567.
15. Vu, Q. D. *et al.* Methods for Segmentation and Classification of Digital Microscopy Tissue Images. *Front. Bioeng. Biotechnol.* **7**, (2019).

16. Horwath, J. P., Zakharov, D. N., Mégret, R. & Stach, E. A. Understanding important features of deep learning models for segmentation of high-resolution transmission electron microscopy images. *Npj Comput. Mater.* **6**, 1–9 (2020).
17. Lugagne, J.-B., Lin, H. & Dunlop, M. J. DeLTA: Automated cell segmentation, tracking, and lineage reconstruction using deep learning. *PLOS Comput. Biol.* **16**, e1007673 (2020).
18. Kromp, F. *et al.* Deep Learning architectures for generalized immunofluorescence based nuclear image segmentation. (2019).
19. Vuola, A. O., Akram, S. U. & Kannala, J. Mask-RCNN and U-net Ensembled for Nuclei Segmentation. *ArXiv190110170 Cs* (2019).
20. Korfhage, N. *et al.* Detection and segmentation of morphologically complex eukaryotic cells in fluorescence microscopy images via feature pyramid fusion. *PLOS Comput. Biol.* **16**, e1008179 (2020).
21. Liu, D. *et al.* Unsupervised Instance Segmentation in Microscopy Images via Panoptic Domain Adaptation and Task Re-Weighting. in *2020 IEEE/CVF Conference on Computer Vision and Pattern Recognition (CVPR)* 4242–4251 (IEEE, 2020). doi:10.1109/CVPR42600.2020.00430.
22. Masubuchi, S. *et al.* Deep-learning-based image segmentation integrated with optical microscopy for automatically searching for two-dimensional materials. *Npj 2D Mater. Appl.* **4**, 1–9 (2020).
23. Deng, J. *et al.* ImageNet: A large-scale hierarchical image database. in *2009 IEEE Conference on Computer Vision and Pattern Recognition* 248–255 (2009). doi:10.1109/CVPR.2009.5206848.
24. Caicedo, J. C. *et al.* Nucleus segmentation across imaging experiments: the 2018 Data Science Bowl. *Nat. Methods* **16**, 1247–1253 (2019).
25. Kromp, F. *et al.* An annotated fluorescence image dataset for training nuclear segmentation methods. *Sci. Data* **7**, 262 (2020).

26. Torr, A., Basaran, D., Sero, J., Rittscher, J. & Sailem, H. DeepSplit: Segmentation of Microscopy Images Using Multi-task Convolutional Networks. in *Medical Image Understanding and Analysis* (eds. Papież, B. W., Namburete, A. I. L., Yaqub, M. & Noble, J. A.) vol. 1248 155–167 (Springer International Publishing, 2020).
27. Schwendy, M., Unger, R. E. & Parekh, S. H. EVICAN—a balanced dataset for algorithm development in cell and nucleus segmentation. *Bioinformatics* **36**, 3863–3870 (2020).
28. Stringer, C., Wang, T., Michaelos, M. & Pachitariu, M. Cellpose: a generalist algorithm for cellular segmentation. *Nat. Methods* 1–7 (2020) doi:10.1038/s41592-020-01018-x.
29. Ljosa, V., Sokolnicki, K. L. & Carpenter, A. E. Annotated high-throughput microscopy image sets for validation. *Nat. Methods* **9**, 637 (2012).
30. Xing, F. *et al.* Towards pixel-to-pixel deep nucleus detection in microscopy images. *BMC Bioinformatics* **20**, 472 (2019).
31. Yang, L. *et al.* NuSeT: A deep learning tool for reliably separating and analyzing crowded cells. *PLOS Comput. Biol.* **16**, e1008193 (2020).
32. Al-Kofahi, Y., Lassoued, W., Lee, W. & Roysam, B. Improved Automatic Detection and Segmentation of Cell Nuclei in Histopathology Images. *IEEE Trans. Biomed. Eng.* **57**, 841–852 (2010).
33. Roth, H. R. *et al.* DeepOrgan: Multi-level Deep Convolutional Networks for Automated Pancreas Segmentation. in *Medical Image Computing and Computer-Assisted Intervention -- MICCAI 2015* (eds. Navab, N., Hornegger, J., Wells, W. M. & Frangi, A.) 556–564 (Springer International Publishing, 2015). doi:10.1007/978-3-319-24553-9_68.
34. Chen, H., Qi, X., Yu, L. & Heng, P.-A. DCAN: Deep Contour-Aware Networks for Accurate Gland Segmentation. *ArXiv160402677 Cs* (2016).
35. Höfener, H. *et al.* Deep learning nuclei detection: A simple approach can deliver state-of-the-art results. *Comput. Med. Imaging Graph.* **70**, 43–52 (2018).

36. Krizhevsky, A., Sutskever, I. & Hinton, G. E. ImageNet Classification with Deep Convolutional Neural Networks. in *Advances in Neural Information Processing Systems 25* (eds. Pereira, F., Burges, C. J. C., Bottou, L. & Weinberger, K. Q.) 1097–1105 (Curran Associates, Inc., 2012).

37. Khadangi, A., Boudier, T. & Rajagopal, V. EM-net: Deep learning for electron microscopy image segmentation. *bioRxiv* 2020.02.03.933127 (2020) doi:10.1101/2020.02.03.933127.

38. Ronneberger, O., Fischer, P. & Brox, T. U-Net: Convolutional Networks for Biomedical Image Segmentation. *ArXiv150504597 Cs* (2015).

39. Raza, S. E. A. *et al.* Micro-Net: A unified model for segmentation of various objects in microscopy images. *Med. Image Anal.* **52**, 160–173 (2019).

40. Shorten, C. & Khoshgoftaar, T. M. A survey on Image Data Augmentation for Deep Learning. *J. Big Data* **6**, 60 (2019).

41. Berryman, S., Matthews, K., Lee, J. H., Duffy, S. P. & Ma, H. Image-based Cell Phenotyping Using Deep Learning. *bioRxiv* 817544 (2019) doi:10.1101/817544.

42. Challen, G. A. & Goodell, M. A. Promiscuous Expression of H2B-GFP Transgene in Hematopoietic Stem Cells. *PLOS ONE* **3**, e2357 (2008).

43. Tumbar, T. *et al.* Defining the epithelial stem cell niche in skin. *Science* **303**, 359–363 (2004).

44. Chalfoun, J. *et al.* FogBank: a single cell segmentation across multiple cell lines and image modalities. *BMC Bioinformatics* **15**, 431 (2014).

45. Stylianidou, S., Brennan, C., Nissen, S. B., Kuwada, N. J. & Wiggins, P. A. SuperSegger: robust image segmentation, analysis and lineage tracking of bacterial cells. *Mol. Microbiol.* **102**, 690–700 (2016).

46. Gurari, D. *et al.* How to Collect Segmentations for Biomedical Images? A Benchmark Evaluating the Performance of Experts, Crowdsourced Non-experts, and Algorithms. *2015 IEEE Winter Conf. Appl. Comput. Vis.* (2015) doi:10.1109/WACV.2015.160.
